# The molecular and metabolic program for adaptation of white adipocytes to cool physiologic temperatures

**DOI:** 10.1101/2020.10.16.342220

**Authors:** Hiroyuki Mori, Colleen E. Dugan, Akira Nishii, Ameena Benchamana, Ziru Li, Thomas S. Cadenhead, Arun K. Das, Charles R. Evans, Katherine A. Overmyer, Steven M. Romanelli, Sydney K. Peterson, Devika P. Bagchi, Callie A. Corsa, Julie Hardij, Brian S. Learman, Mahmoud El Azzouny, Ken Inoki, Ormond A. MacDougald

## Abstract

Although visceral adipocytes located within the body’s central core are maintained at ~37°C, adipocytes within bone marrow, subcutaneous, and dermal depots are found primarily within the peripheral shell, and generally exist at cooler temperatures. Responses of brown and beige/brite adipocytes to cold stress are well-studied; however, comparatively little is known about mechanisms by white adipocytes adapt to temperatures below 37°C. Here we report that adaptation of cultured adipocytes to 31°C, the temperature at which distal marrow adipose tissues and subcutaneous adipose tissues often reside, induces extensive changes in gene expression, increased anabolic and catabolic lipid metabolism, and elevated oxygen consumption with reduced reliance on glucose and preferential use of pyruvate, glutamine and fatty acids as energy sources. Cool temperatures up-regulate stearoyl-CoA desaturase-1 expression and monounsaturated lipid levels in cultured adipocytes and distal bone marrow adipose tissues, and stearoyl-CoA desaturase-1 activity is required for acquisition of maximal oxygen consumption at 31°C.

## INTRODUCTION

Survival of euthermic animals is dependent on tight regulation of body temperature and cellular function in environmental conditions below thermoneutrality. Although mammals have developed a complex system to defend core body temperature within a narrow range, the exterior and extremities remain much cooler. Whereas the body core is maintained at a relatively constant ~37°C across a broad range of environmental conditions, the body shell, which includes peripheral extremities and subcutaneous regions of the trunk, is characterized by a wider spectrum of temperatures. For almost a century, scientists have known that temperature gradients exist across the body, from central to distal, and superficial to deeper regions (1). For example, subcutaneous tissues in the human calf are generally ~32°C but can dip to below 30°C in the cold and increase to 37°C when it is hot or during exercise (1, 2). Even subcutaneous tissues of the chest and back are generally 2°C below core temperature and function in a more dynamic range (32-37°C) than cells located deeper within the body (2).

Temperature is detected by sensory neurons within skin, core, and brain, which transmit signals to the central thermoregulatory network. After signal integration, specific hypothalamic neurons regulate body temperature through a combination of behavioral and physiological mechanisms (3, 4). Physiological responses to cold include vasoconstriction within skin to limit cooling, as well as shivering and adaptive thermogenesis to generate heat. The canonical adaptive thermogenesis pathway involves activation of sympathetic drive to increase brown adipose tissue metabolism and UCP1-dependent uncoupling of the mitochondrial proton gradient from ATP synthesis (5, 6). Activation of adaptive thermogenesis may be an intrinsic cellular response in some contexts - moving 3T3-F442A adipocytes from 37°C to cool temperatures (27-33°C) rapidly induces expression of UCP1 and uncoupled oxygen consumption (7). In addition, recent work has uncovered non-canonical mechanisms for heat generation in beige/brite adipocytes, including a creatine kinase futile cycle (8, 9) and ATP-dependent calcium cycling mediated by SERCA2b (10, 11). Whereas the response of brown and beige/brite adipocytes to cold stress has been extensively studied, comparatively little is known about the program by which white adipocytes adapt to cool temperatures.

White adipocytes are distributed throughout the body in discrete depots and the intrinsic cellular and metabolic properties of different populations are shaped by the specific niches in which they reside (12, 13). Whereas visceral white adipocytes are found within the body’s core, other populations, including subcutaneous, marrow and dermal adipocytes, primarily exist in temperatures well below 37°C. Despite this, studies within the field have largely neglected environmental temperature as a variable when considering the molecular and functional characteristics of white adipocytes. Although exposure of white adipocytes to cooler temperatures does not induce known adaptive thermogenesis pathways, hints from historical literature suggest that cooler adipose tissue temperatures are associated with greater lipid unsaturation (14). Tavassoli et al and our own work have found similar correlations in rabbits, rodents and humans, with distal marrow adipocytes containing a larger proportion of unsaturated lipids and proximal sites (15, 16). How cool temperature-adapted cells, such as subcutaneous and distal marrow adipocytes, are functionally different from warmer counterparts is unknown.

To explore adaptation of white adipocytes to changing environmental temperature, we exposed cultured adipocytes to a “cool” temperature of 31°C, the temperature at which distal marrow and subcutaneous adipose tissues exist when animals and humans are at a room temperature of 22°C (1, 2, 17). We found that exposure of cultured adipocytes or adipose tissues to cool temperatures increases expression of stearoyl-CoA desaturase-1 (SCD1) and levels of monounsaturated lipids within triacylglycerols (TAG). SCD1 is highly expressed in adipocytes and converts saturated lipids such as palmitoyl-CoA to monounsaturated lipids such as palmitoleic acid. Culturing adipocytes at 31°C causes extensive changes in gene expression, increased anabolic and catabolic lipid metabolism, and elevated oxygen consumption with reduced reliance on glucose and preferential use of pyruvate, glutamine and fatty acids as energy sources. In addition to expansion of mitochondria and peroxisomes, we observed up-regulation of proteins involved in β-oxidation and complex I, II and III of the electron transport chain. Furthermore, SCD1 activity is required for the increase in maximal oxygen consumption and spare respiratory capacity observed during cool adaptation. Thus, the molecular and metabolic program for adaptation of white adipocytes to cooler temperatures is cell-autonomous and distinct from canonical thermogenesis.

## RESULTS

### Cool environmental temperatures induce SCD1 expression and lipid desaturation

We previously reported that isolated bone marrow adipocytes (BMAds) from the distal tibia (dTib) of mice have elevated *Scd1* expression and higher levels of monounsaturated lipids compared to those found in the proximal tibia or within white adipose depots (15). We hypothesized that this increase in lipid desaturation occurs because distal BMAds exist on the cool end of temperature gradients present in rodents housed at 22°C. Indeed, in rodents housed at 22°C, temperatures across caudal vertebrae (CV) range from 38°C at CV1 and CV2 to 32°C by CV9 (17). To test this hypothesis, we housed rats at room temperature (22°C) or thermoneutrality (29°C) from birth to 11 weeks of age. Lipidomic analyses revealed decreased levels of saturated fatty acids (myristic, palmitic and stearic acids; shades of blue) and increased proportions of unsaturated fatty acids, particularly oleic acid, esterified within TAG isolated from distal bone marrow adipose tissue (BMAT) depots of rats housed at 22°C (**Figure 1a**). Interestingly, we did not detect temperature-dependent alterations in phospholipids (**Figure S1a**). We then focused our analyses on SCD, an endoplasmic reticulum enzyme that catalyzes formation of monounsaturated fatty acids; specifically, it converts Coenzyme A-derivatives of myristic (C14:0), palmitic (C16:0) and stearic (C18:0) acids to myristoleic (C14:1, n-5), palmitoleic (C16:1, n-7) and oleic (C18:1, n-9) acids, respectively. Consistent with the lipidomic profile, SCD1 expression is also regulated by temperature, with considerably more SCD1 protein observed in CV and dTib BMAT depots of rats housed at 22°C compared to 29°C (**Figure 1b**).

**Figure 1.**
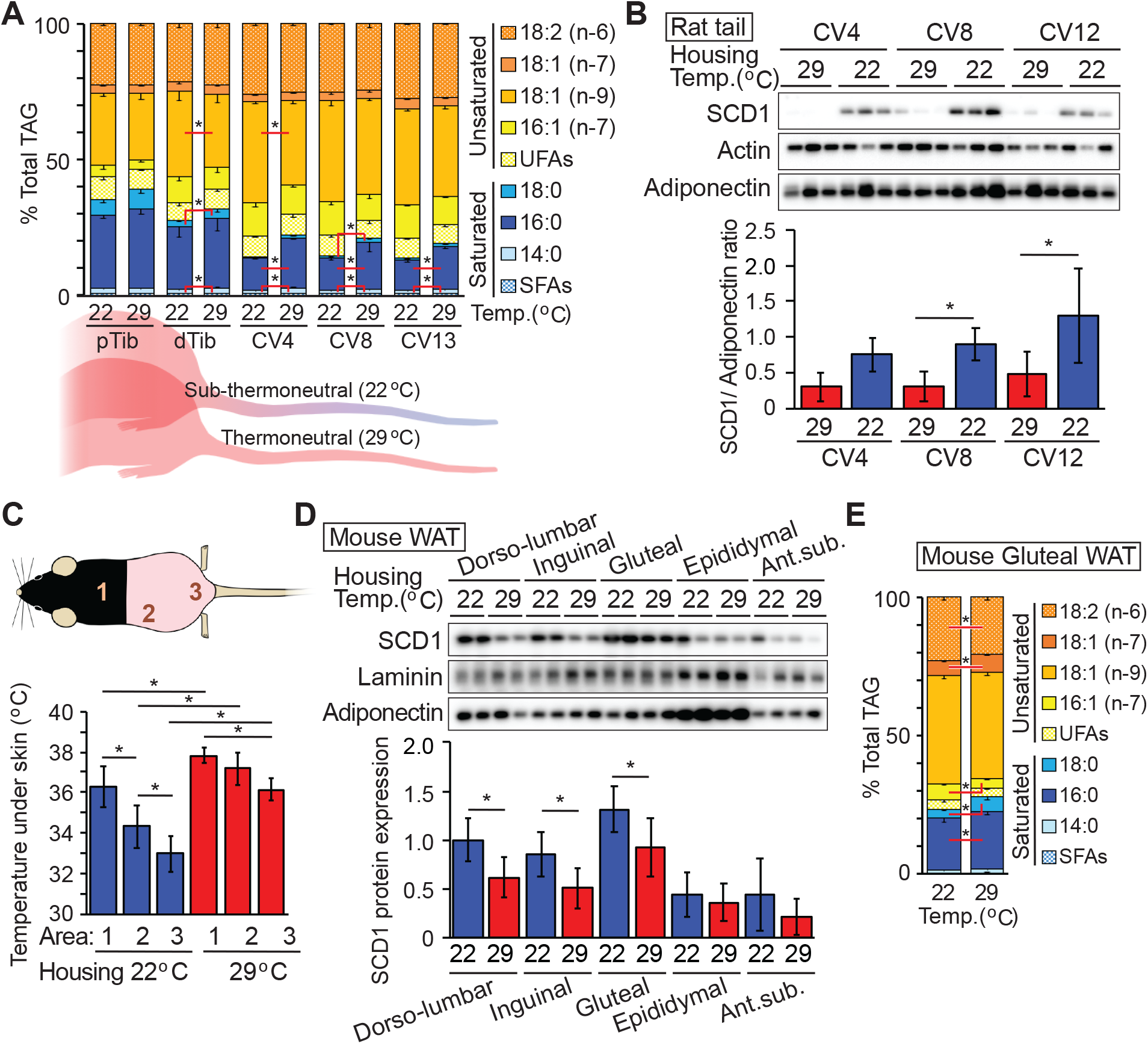
Lipid unsaturation and SCD1 expression are induced by cool environmental temperatures. (**a**) Increased proportion of unsaturated lipid in BMAT TAG of distal tibia and caudal vertebra of rats housed at 22°C. Rats were housed from birth to 11 weeks of age at standard room temperature (22°C) or thermoneutrality (29 °C). Lipid composition of TAG for proximal (pTib) and distal tibia (dTib), and indicated caudal vertebra (CV) were determined by gas chromatography. (*n* = 6 except CV4 and 13; *n* = 3). (**b**) Expression of SCD1 is elevated in CV from animals housed at 22°C compared to 29°C. SCD1 protein levels were normalized to adiponectin (*n* = 7-9). (**c**-**e**) Mice were housed from birth to 13 weeks of age at 22°C or 29°C. After weaning, posterior hair was removed. (**c**) The subdermal temperature of mice housed at each temperature (*n* = 7, 8). (**d**) Elevated SCD1 expression in subcutaneous WAT depots of mice at 22 °C. SCD1 protein expression was normalized to geometric mean value of the following proteins; adiponectin, perilipin, HSP70, and laminin (*n* = 7 or 8). (**e**) Increased proportion of unsaturated lipid in TAG of gluteal WAT of mice housed at 22°C (*n* = 5). Data are presented as mean ± s.d. \**p*<0.05.

To explore if changes in gene expression induced by cool temperatures are also observed in white adipose tissues (WAT), similar experiments were performed using mice exposed to cool environmental temperatures. In a subset of mice, posterior hair was removed to reduce insulation. In mice housed at 22°C, subcutaneous temperatures were found to be ~33-34°C, approximately 3°C lower than in mice housed at 29°C (**Figure 1c**). Consistent with this reduced temperature, SCD1 expression is higher in posterior subcutaneous depots (dorsolumbar, inguinal and gluteal WAT) of mice housed at 22°C; in contrast, SCD1 is not altered in anterior subcutaneous or visceral WAT depots, where tissue temperatures are maintained at ~37°C (**Figure 1d** and **Figure S1b**). Expression of adiponectin is not altered by environmental temperature and serves as a loading control (**Figure 1d** and **Figure S1c**). Consistent with the SCD1 expression, we observed decreased proportions of saturated fatty acids (palmitic and stearic acids) and increased palmitoleic and vaccenic acids within TAG of gluteal WAT of mice housed at 22°C (**Figure 1e**). These data indicate that monounsaturated lipid content is increased by exposure to cooler temperatures in both BMAT and posterior subcutaneous WAT.

### Effects of cool temperatures on increased SCD1 expression and lipid monounsaturation are cell-autonomous

To study the regulation of lipid desaturation and SCD1 expression by temperature more mechanistically, we established a cell culture system in which we could evaluate how white adipocytes adapt to cool temperatures. We chose to use mesenchymal stem cells (MSCs) because they can be isolated directly from transgenic mouse models of interest and are highly adipogenic (18). A subpopulation of adipose tissues, such as subcutaneous WAT, distal BMAT, and dermal WAT, exist at temperatures as low as 31°C when animals are housed at 22°C (2, 17). Thus, we chose 31°C as the “cool” temperature to mimic the physiological conditions under which adipocytes reside *in vivo*. MSCs were cultured at 37°C prior to and for the first four days of adipogenesis, after which cells were moved to 31°C for 12 days. Exposure to 31°C does not influence the late stages of adipogenesis, as assessed by cell morphology and expression of adipocyte markers, such as PPARγ, adiponectin, and FABP4 (**Figure S2a** and **S2b**). Importantly, beige adipocyte markers, including *Ucp1*, *Fgf21*, and *Pgc1a*, are not induced in white adipocytes cultured at 31°C (**Figure S2b** and **S2c**). Consistent with our *in vivo* results (**Figure 1**), expression of SCD1 mRNA and protein is increased in adipocytes cultured at temperatures below 37°C (**Figure 2a** and **2b**). Indeed, changes in SCD1 expression are observed with shifts as small as 2°C, from 37°C to 35°C, but particularly with the shift to 33°C. Effects of culturing cells at 31°C on SCD1 protein emerge as early as 12 hours after exposure, with further increases observed until 96 hours (**Figure 2d**); in contrast, effects on *Scd1* mRNA are more subtle, only becoming significant after exposure to 31°C for 96 hours (**Figure 2c**). As expected, adipocytes cultured at 31°C for 11 days have increased unsaturation within TAG lipids, characterized by a significant switch from palmitic to palmitoleic acid (**Figure 2e**), with milder changes observed in PL composition. Similarly, a higher proportion of double bonds are observed in TAG following cool adaptation (**Figure 2f**). Taken together, these data indicate that effects of cool temperature on induction of SCD1 and increased lipid monounsaturation are cell-autonomous.

**Figure 2.**
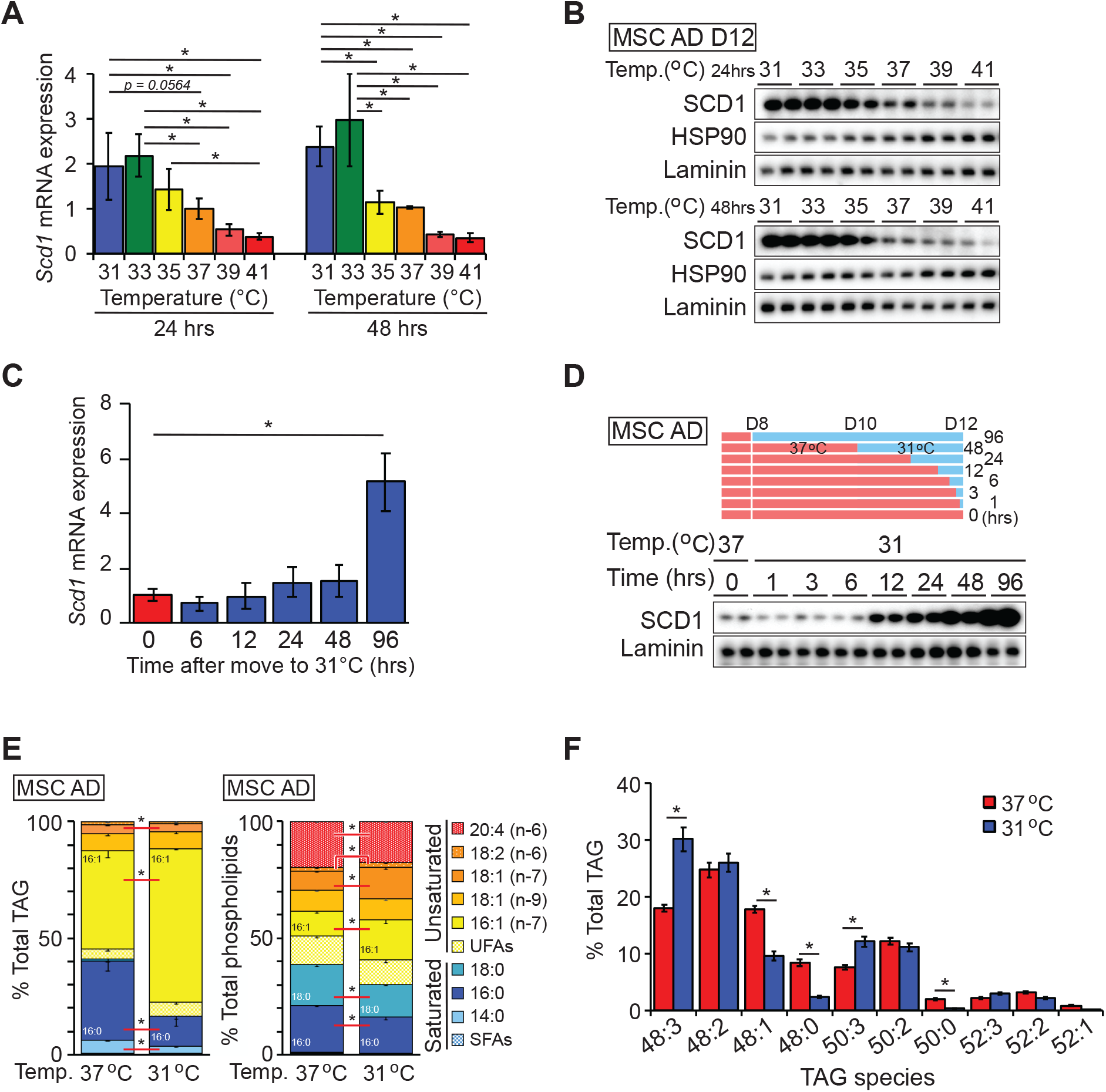
Adaptation of cultured adipocytes to cool temperatures induces expression of SCD1 and proportion of unsaturated lipids in TAG and phospholipid fractions. Cultured adipocytes were incubated for 24 or 48 hours at indicated temperatures before collecting samples. (**a-b**) Expression of SCD1 mRNA (**a**) and protein (**b**) were evaluated by qPCR and immunoblot respectively. *Scd1* mRNA was normalized to the geometric mean value of *Hprt*, *Tbp*, *Gapdh*, *Rpl32*, *Rpl13a*, *B2m*, and *Rn18s,* and is expressed as fold change relative to 37°C control (*n* = 4). (**c-d**) SCD1 mRNA (**c**) and protein (**d**) expression in adipocytes cultured at 31°C for indicated times. SCD1 mRNA was normalized to *Hprt* mRNA and is expressed as fold change relative to 0 h control (*n* = 4). (**e**) Lipid composition of TAG and phospholipid fractions in adipocytes adapted to 31°C for 11 days (*n* = 6) with cool adaption. (**f**) TAG species in adipocytes adapted to 31°C for 11 days (*n* = 4). Data are presented as mean ± s.d. \**p*<0.05.

### Exposure of adipocytes to 31°C activates an extensive genetic program of adaptation

To evaluate the genetic program underlying cool adaptation, we performed RNA-seq on mature adipocytes exposed to 31°C for 0, 1 or 12 days. Twelve days of cool adaptation caused rapid and large-scale changes in gene expression, with up-regulation of 1,872 genes and suppression of 2,511 genes (**Figure 3a** and **Figure S3**). Following either 1 or 12 days of cool exposure, we did not observe regulation of common adipocyte genes such as *Cebpa*, *Creb*, *Klfs*, *Glut4*, *Plin*, *Lpl*, and *Bscl* (**Supplemental data**), consistent with lack of obvious morphological differences. Markers of brown and beige adipocytes (e.g. *Ucp1*, *Ppargc1a*, *Eva1*, *Tnfsf9*, *Cox8b*, *Fgf21*) (19) are also not induced in this context (**Supplemental data**). Similarly, genes involved in the creatine kinase cycle (*Slc6a8*, *Gamt*, *Ckmt1*, *Ckmt2*) (8) or the SERCA2b-dependent futile cycle (*ATP2a2*) (10) are not altered by exposure to 31°C for 1 or 12 days (**Supplemental data**). Further, evaluation of genes involved in metabolism suggests that cool exposure results in altered amino acid and glutathione metabolism, suppressed glycolysis/gluconeogenesis and pentose shunt pathways, and elevated lipid degradation (**Figure 3b**). Gene set enrichment analysis (GSEA) revealed that 12 days of 31°C exposure significantly induces genes involved in oxidative metabolism, as well as cell cycle regulation, including E2F and Myc targets, mitotic spindle, and G2M checkpoints. GSEA also predicts that cool adaptation suppresses glycolysis, along with impaired response to interferon gamma and hypoxia (**Figure 3c**). Thus, exposure of primary cultured adipocytes to 31°C for 1 or 12 days results in extensive transcriptome changes that suggest that non-glucose energy sources support elevated oxygen consumption.

**Figure 3.**
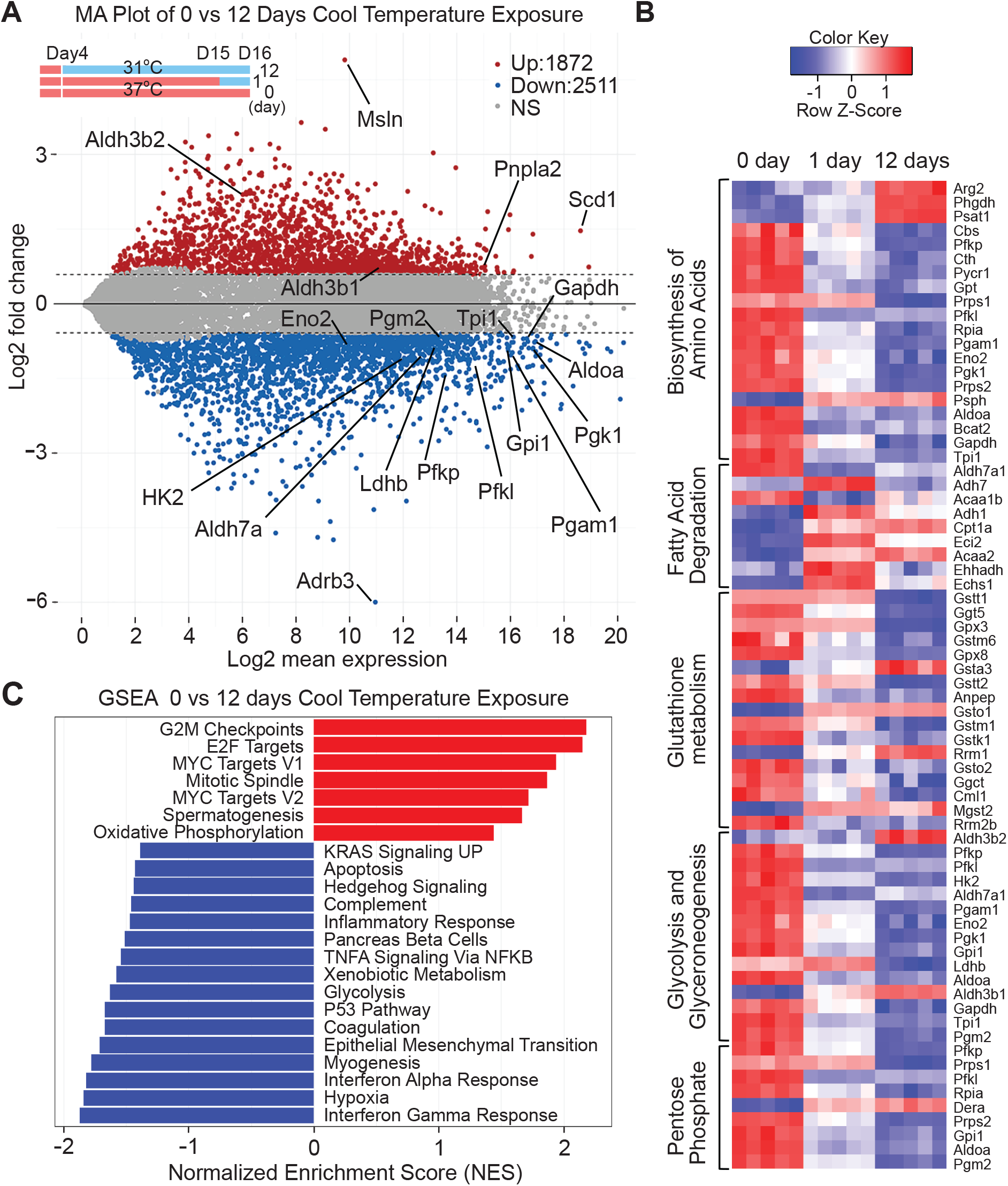
Exposure of adipocytes to 31°C activates an extensive genetic program of adaptation. RNA from mature adipocytes at day 0, 1 and 12 of cool adaptation was purified and subjected to RNA-Seq analyses (*n* = 5 per time point). (**a**) MA plot showing the log2-mean expression versus log2-fold change of mRNA transcript expression in 12-day cool exposed MSC adipocytes compared to day 0. Each dot represents a gene. Twelve days of cool temperature exposure induced 1872 genes (red) and suppressed 2511 genes (blue). Significance was defined by a FDR < 0.05 and absolute fold change > 1.5. (**b**) Heat map of select metabolic KEGG pathways significantly enriched by 1 or 12 days of cool temperature from iPathwayGuide pathway impact analysis. Enrichment was defined by a nominal p value < 0.05 and FDR < 0.25. (**c**) Normalized score plots of enriched pathways from preranked Gene Set Enrichment Analysis comparing RNA expression in mature adipocytes exposed to 31°C for 0 or 12 days. Enriched pathways were defined by a nominal p value < 0.05 and FDR < 0.05. P-values were calculated from GSEA with 1,000 permutations. Data shown is from one experiment.

### Adipocytes adapted to 31°C have increased oxygen consumption rates (OCR) and altered nutrient selection and utilization

To test predictions from RNAseq analyses, we next evaluated OCR of cool-adapted adipocytes using a Seahorse XF Analyzer (**Figure 3c**). To control for potential effects of short-term temperature changes on oxidative processes, both cool-exposed and control adipocytes were analyzed at both 37°C and 31°C. When assays were performed at either 37°C or 31°C, cool adaptation increased basal OCR ~2.5-fold (**Figure 4a** and **4b**). Whereas the mitochondrial capacity for respiration was elevated when cool-adapted adipocytes were evaluated with FCCP at 31°C (**Figure 4a**), this difference was masked at 37°C by the dramatic increase to near-maximal rates of oxidative metabolism (**Figure 4b**). Of note, adipocytes cultured at 37°C also exhibit reduced rates of metabolism when assayed at 31°C rather than at 37°C. In cool-adapted adipocytes assayed at 31°C, we observed elevated non-mitochondrial OCR following treatment with rotenone/antimycin, which is generally attributed to other cellular oxidative reactions, including peroxisomal respiration; however, this difference was not observed in cells assayed at 37°C.

**Figure 4.**
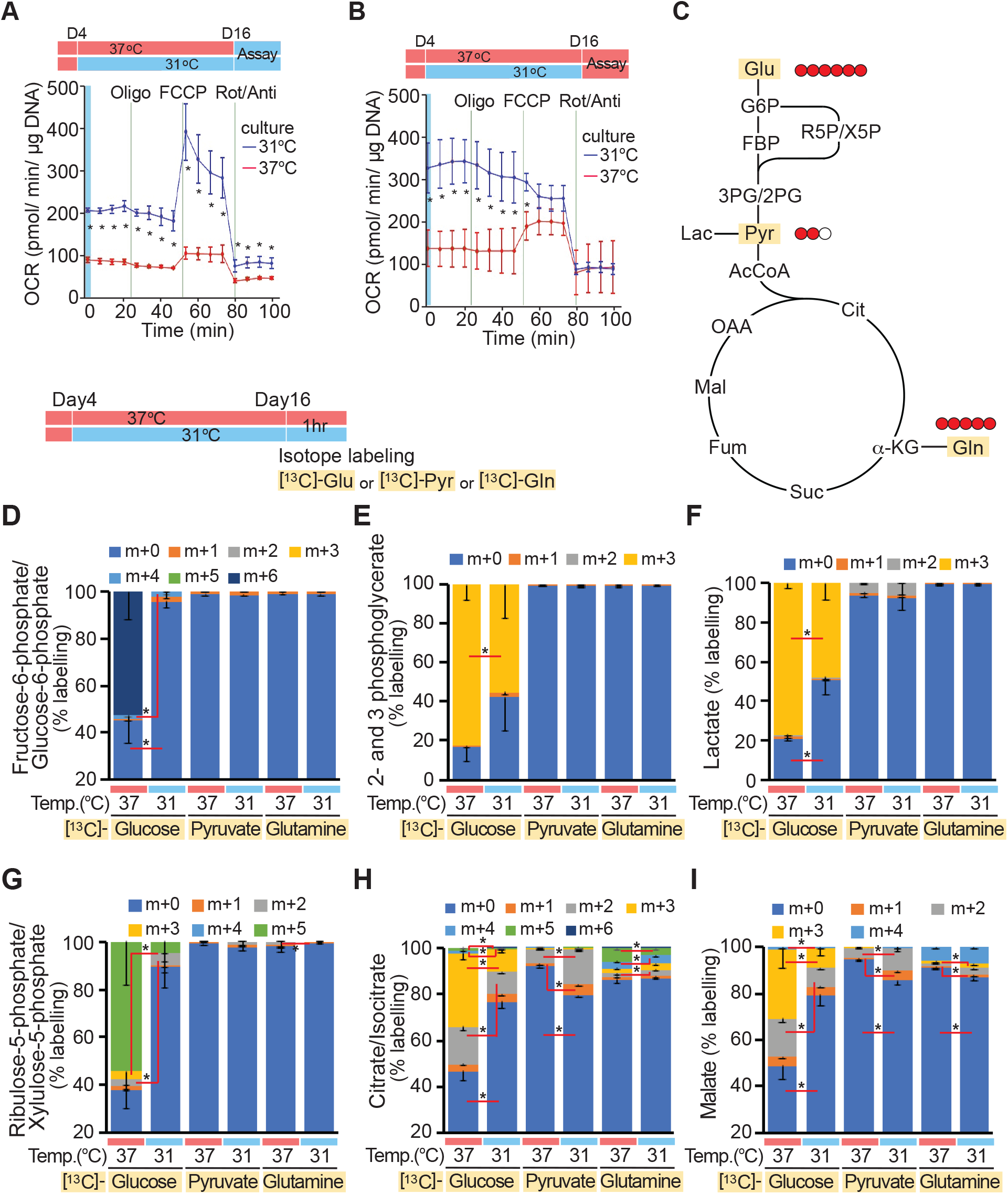
Adipocytes adapted to cooler temperatures have increased OCR, and preferentially consume pyruvate and glutamine over glucose. (**a-b**) Seahorse XF96 Cell Culture Microplates were cut in half. Four days after adipocyte differentiation at 37°C, adipocytes on one half of the plate was cultured at 31°C and the other half at 37°C for 12 days. After rejoining the plate, OCR was evaluated either at 31°C (**a**) or 37°C (**b**) (*n* = 4). Oligo: oligomycin, Rot/Anti: rotenone and antimycin. Data shown is representative of at least 3 independent experiments. (**c**-**i**) Differentiated adipocytes cultured at either 37°C or 31°C for 12 days were incubated with [^13^C]-glucose, [^13^C]-pyruvate or [^13^C]-glutamine for 1 h. Substrate concentrations in media were kept constant except for the substitution of [^13^C]-metabolites (5 mM glucose, 0.2 mM pyruvate and 1 mM glutamine). Color code indicates [^13^C]-labeling on 0 through 6 carbons (*n* = 3). Values are mean ± s.d. \**p* < 0.05. Data shown is from one experiment.

We next evaluated which substrates are utilized for increased oxygen consumption in adipocytes adapted to 31°C. RNA-seq analyses suggested that flux through glycolysis and the pentose phosphate shunt would be suppressed with cool adaptation (**Figure 3b** and **3c**); thus, we performed metabolic flux analyses using isotope-labeled glucose, pyruvate and glutamine as carbon tracers. Adipocytes were cultured at 31°C or 37°C for 12 days and then incubated for 1 hour with [^13^C]-glucose, [^13^C]-pyruvate or [^13^C]-glutamine (**Figure 4c**). Cool-adapted adipocytes have reduced flux of glucose through glycolysis, as assessed by [^13^C]-glucose labeling of fructose-6-phosphate/glucose-6-phosphate (**Figure 4d**), 2- and 3-phosphoglycerate (**Figure 4e**) and lactate (**Figure 4f**), and through the pentose phosphate shunt, as assessed by [^13^C]-glucose labeling of ribulose-5-phosphate and xylulose-5-phosphate (**Figure 4g**). There is also reduced flow of [^13^C]-glucose into tricarboxylic acid cycle intermediates, such as citrate/isocitrate (**Figure 4h**) and malate (**Figure 4i**). In contrast, a significant proportion of labeled carbons from [^13^C]-pyruvate and [^13^C]-glutamine are incorporated into citrate/isocitrate and malate (**Figure 4h, 4i**). Thus, adaptation to 31°C increases basal OCR, and both transcriptomics and metabolomic flux analyses point to diminished reliance on glucose for oxidative metabolism. Further, these results suggest that adipocytes adapted to cooler temperatures preferentially consume pyruvate and glutamine over glucose.

### Fatty acid oxidation is increased in cultured adipocytes adapted to cool environmental temperatures

Adipocytes adapted to 31°C shift carbon sources away from glucose towards glutamine and pyruvate (**Figure 4**). Although amino acids are anaplerotic substrates in adipocytes (20), glucose and fatty acids are considered to be primary carbon sources for their energy needs. To explore further how cool adaptation influences adipocyte lipid metabolism, we measured β-oxidation in adipocytes exposed to 31°C for different lengths of time. We observed that release of tritiated water from adipocytes treated with [^3^H]-palmitic acid is elevated after six days of exposure to 31°C, and increases further with 8 and 12 days of cool adaptation (**Figure 5a**). Increased β-oxidation in cool-adapted adipocytes is also observed when labeled oleic acid is used as a substrate (**Figure 5b**). Interestingly, oxidation of octanoic acid (C8:0; **Figure S4a**) is only slightly increased with cool adaptation, suggesting that mitochondrial uptake of fatty acids is highly regulated by cool-adaptation. In this regard, proteins involved in both uptake and β-oxidation of fatty acids, including carnitine palmitoyltransferase 1α (CPT1α), medium-chain 3-ketoacyl-CoA thiolase (MCKAT), and mitochondrial trifunctional protein beta subunit (MTPβ), are up-regulated by cool adaptation; these effects are observed in several cell models, including cultured adipocytes differentiated from MSCs (**Figure 5c**) or stromal-vascular cells (**Figure S4b, S4c**), and primary white adipocytes isolated by collagenase digestion of WAT (**Figure 5d**). Pharmacological inhibition of mitochondrial fatty acid uptake (**Figure 5a**) or OCR (**Figure 4a, 4b**) indicates that there is also a significant non-mitochondrial component involved in cool adaptation. This may be due to elevated peroxisomal α - or β-oxidation, as evidenced by increased expression of peroxisome markers, such as PMP70 and PEX5 (**Figure 5c**). Since peroxisomal oxidation of fatty acids is not coupled to ATP synthesis, it may contribute to heat generation in cool-adapted adipocytes.

**Figure 5.**
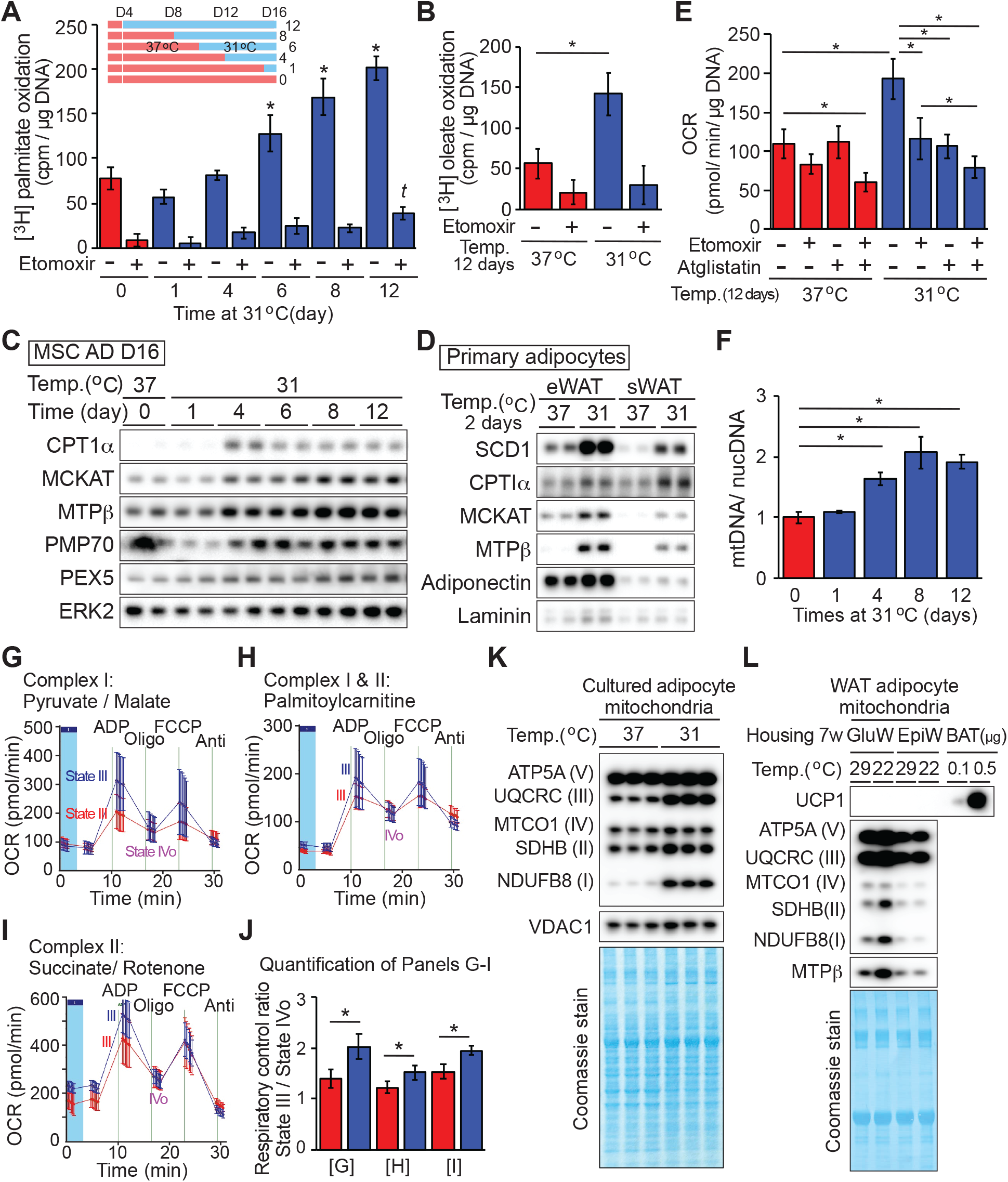
Cool adaptation of adipocytes stimulates β-oxidation of fatty acids and increases number and functioning of mitochondria. (**a**-**b**) Adipocytes at indicated days of cool temperature exposure (**a**; *n* = 6) or for 12 days (**b**; *n* = 5) were incubated with tritiated palmitate (**a**) or oleate (**b**) for 3 h +/− 40 μM etomoxir. * significantly different from day 0 control at 37°C; *p* < 0.05). *t* indicates a difference from day 0 etomoxir treatment. (**c**) Cool adaptation increases enzymes involved in fatty acid oxidation and peroxisomal protein markers. (**d**) Immunoblot analyses of primary adipocytes isolated from epididymal white adipose tissue (eWAT) or subcutaneous gluteal WAT (sWAT) were cultured floating at either 37°C or 31°C for 2 days. (**e**) Elevated basal OCR of adipocytes at 31°C is partially dependent on oxidation of endogenous fatty acids. MSC-derived adipocytes cultured at 31°C or 37°C for 12 days were treated with vehicle, 20 μM etomoxir and/or 20 μM of atglistatin for 2 hr prior and during assay of OCR (*n* = 8-10). (**f**) Four days exposure at 31°C stimulates an increase in mitochondrial number as assessed by qPCR for two mitochondrial regions (CytoB, Cox-TMS) and two nuclear genes (glucagon, β-globin; *n* = 3). (**g**-**i**) Isolated mitochondria from adipocytes adapted to 31°C have improved functioning of complex I and II. Mitochondria were incubated with media containing pyruvate/ malate for NADH-linked (Complex I) substrates (**g**), Palmitoylcarnitine/malate for Complex I and II (**H**), and succinate/rotenone for Complex II (**i**). (**j**) Calculated respiratory control ratios (RCR) of mitochondria from panel (**G**) to (**I**), (*n* = 3 per group), values are mean ± s.d. \**p* < 0.05. (**k**) Isolated mitochondria were lysed and proteins were quantified. OXPHOS proteins were evaluated by immunoblot. VDAC1 is used as a loading control and Coomassie blue staining of the membrane shows other mitochondrial proteins. (**l**) Mitochondria were isolated from floating adipocytes following collagenase digestion of gluteal or epididymal WAT from 5 mice housed at either at 29°C or 22°C for 7 weeks. OXPHOS proteins were evaluated and Coomassie staining was performed. Data are presented as mean ± s.d. \**p* < 0.05. Data shown is representative of at least 3 independent experiments.

These results led us to investigate further the relationship between fatty acid utilization and increased OCR in adipocytes cultured at 31°C. Pharmacologic inhibition of endogenous fatty acid uptake into mitochondria with the CPT1 inhibitor, etomoxir, decreases basal OCR of cool-adapted adipocytes by ~50%, whereas the OCR of adipocytes cultured at 37°C is not affected (**Figure 5e**). Since the assay media does not contain exogenous fatty acids, treatment with etomoxir exclusively blocks entry of endogenous fatty acids into the mitochondria. Similarly, treatment of cool-adapted adipocytes with the ATGL-specific inhibitor aglistatin blocks lipolysis, thereby decreasing the endogenous fatty acid supply and suppressing basal OCR of cool-adapted adipocytes (**Figure 5e**). Although combined treatment with etomoxir and atglistatin suppresses basal OCR in cells cultured at either 31°C or 37°C, we observed a greater degree of suppression in cool-adapted adipocytes, suggesting that these cells rely more heavily on endogenous fatty acids for energy.

### Isolated mitochondria from adipocytes adapted to 31°C have improved complex I and II function

Increased OCR and fatty acid oxidation in cool-adapted adipocytes led us to next investigate effects on mitochondrial number and function within these cells. Measurement of mitochondrial DNA (mtDNA) relative to nuclear DNA (ncDNA) by qPCR demonstrated that mitochondrial number doubles by eight days of cool adaptation (**Figure 5f**). Furthermore, mRNA expression of most mitochondrial complex subunits is also up-regulated following exposure to 31°C (**Figure S4d**). This mitochondrial expansion is not accompanied by increased expression of *Ppargc1a*, *Tfam*, *Nrf1*, or *Ucp1* (**Supplemental data, RNA-Seq**), although up-regulation of mitochondrial proteins specifically involved in the β-oxidation pathway is observed (**Figure 5c** and **5d**). Although elevated capacity for cellular OCR and fatty acid oxidation with adaptation to 31°C can be explained, in part, by an increase in mitochondrial number (**Figure 5f**), we also evaluated functional characteristics of isolated mitochondria. Mitochondrial respiratory control encapsulates the main function of mitochondria: the ability to idle at a low rate and respond to ADP to generate large quantities of ATP. Interestingly, we found that ADP-stimulated respiration (State III) is higher in mitochondria from adipocytes cultured at 31°C compared to those cultured at 37°C (**Figure 5g-5j**). These results indicate that mitochondria of cool-adapted adipocytes have enhanced function of both complex I (malate plus pyruvate) (**Figure 5g, 5j**) and complex II (succinate plus rotenone) (**Figure 5i, 5j**). These results also hold true when mitochondria are treated with palmitoylcarnitine to test complex I and II together (**Figure 5h, 5j**). No significant changes in non-ADP-stimulated respiration (state IVo), were observed between the two groups following inhibition of ATP synthase by oligomycin. These data suggest that cool-adapted adipocyte mitochondria have a high capacity for substrate oxidation and ATP turnover. Immunoblot analyses of oxidative phosphorylation (OXPHOS) complexes revealed that NDUFB8 of complex I, SDHB of complex II, and UQCRC2 of complex III are increased disproportionately when compared to other mitochondrial proteins evaluated (**Figure 5k**).

Next, to examine whether expression of specific OXPHOS proteins is regulated by temperature *in vivo*, mice were housed at 29°C or 22°C for seven weeks. Mitochondria were then isolated from floating adipocytes following collagenase digestion of subcutaneous gluteal WAT or visceral epididymal WAT. Immunoblot analyses revealed that expression of complex I (NDUFB8) and complex II (SDHB) proteins are elevated in mitochondria isolated from gluteal WAT of mice housed at 22°C (**Figure 5l**), whereas no differences are observed in other complex proteins. These results suggest that in cool-adapted adipocytes, increased oxidative metabolism is due not only due to increased mitochondrial number, but also to formation of more high-functioning, ‘healthy’ mitochondria with improved respiratory capacity. Of note, elevated uncoupled mitochondrial respiration in cool-adapted adipocytes is not due to canonical adaptive thermogenesis, since *Ucp1* mRNA expression is unchanged (**Supplemental data, RNA-Seq**) and protein levels are undetectable in both cultured and primary white adipocytes (**Figure S5a, S5b**). Furthermore, increased OCR in response to cool exposure occurs at similar levels in wildtype and UCP1-deficient adipocytes (**Figure S5d**), with comparable induction of proteins such as SCD1, FASN and CPT1α (**Figure S5c**).

### Adaptation to 31°C for 12 days down-regulates basal and stimulated TAG hydrolysis, but does not limit total lipolytic capacity

RNA-seq analyses identified *Adrb3* as one of the most profoundly suppressed adipocyte genes with cool exposure (**Figure 3a**). *Adrb3* encodes the β3-adrenergic receptor (β3-AR), which is predominantly expressed in rodent adipocytes (21). Activation of the β-AR leads to the activation of protein kinase A, which phosphorylate perilipin and hormone-sensitive lipase (HSL) to promote lipolysis. Thus, we next investigated effects of cool temperatures on basal and stimulated lipolysis. Adipocytes exposed to 31°C or 37°C for 12 days were treated with vehicle, forskolin, or the β3-agonist CL-316,243 (CL) for 6 hrs. Lipolysis was evaluated by measurement of non-esterified fatty acids (NEFA) and glycerol released into media. As expected, adipocytes cultured at 37°C respond robustly to both forskolin and CL by increasing release of glycerol and NEFA (**Figure 5a, 5b**). In contrast, cool-adapted adipocytes demonstrate significantly reduced lipolysis at baseline, and following stimulation with CL. Adipocytes cultured at 31°C and 37°C respond similarly to forskolin treatment, suggesting that reduced secretion of lipolytic products from cool-exposed adipocytes at baseline or in response to CL is not due to decreased lipolytic capacity (**Figure 6a, 6b**).

**Figure 6.**
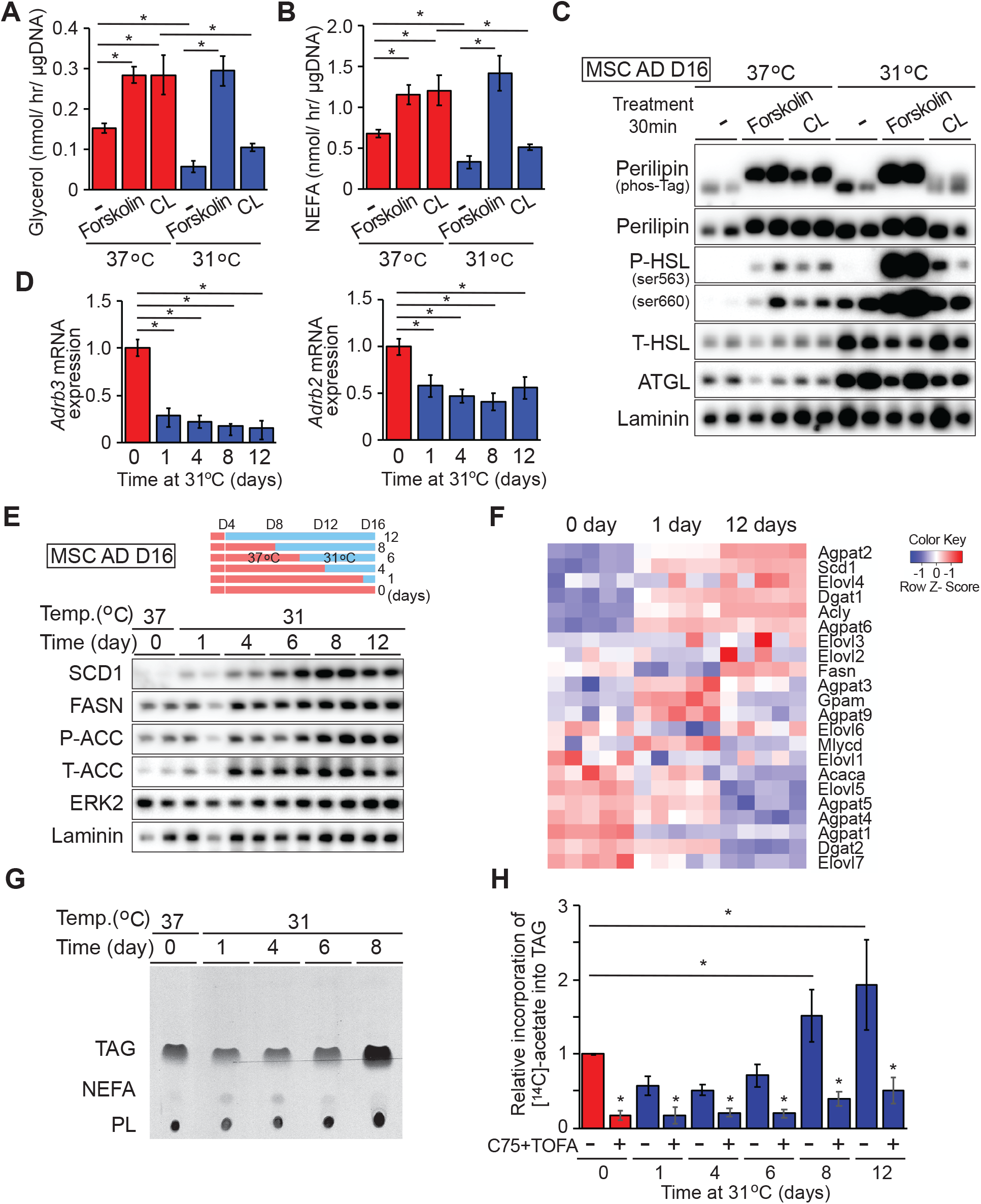
Adaptation to cool temperature decreases basal and stimulated lipolysis and increases *de novo* lipogenesis and TAG synthesis. (**a**-**b**) Adipocytes were treated with 10 μM forskolin or 2 μM CL-316,243 (CL) for 6 hours and secretion of glycerol (**a**) and NEFA (**b**) measured (*n* = 4). (**c**) Cool adaptation increases expression of lipases, but decreases phosphorylation of perilipin and HSL in response to CL-316,243. Adipocytes were treated with vehicle, 10 μM forskolin or 2 μM CL for 30 min, and lysed for immunoblot analyses. (**d**). Rapid reduction of β3- and β2-adrenergic receptor mRNAs after exposure to 31°C (*n* = 6 per time point). Gene expression was normalized to geometric mean of *Hprt*, *Tbp*, *Ppia,* and is expressed relative to 37°C control (*n* = 4). (**e**) Expression of *de novo* lipogenesis-related proteins in adipocytes cultured at 31°C. (**f**) Relative expression of *de novo* lipogenesis-related mRNAs after 0, 1 or 12 days of 31°C. Genes were manually curated since *de novo* lipogenesis is not captured specifically by iPathwayGuide or GSEA. Cool adaptation for 8 days increases incorporation of [^14^C]-acetate into TAG and phospholipids. (**g**) [^14^C]-acetate was added to media of adipocytes cultured at 31°C. Metabolites were separated by thin layer chromatography, and [^14^C] was detected by autoradiography. Addition of 10 μM TOFA and 50 μM of C75 was for 3 h before and throughout the lipogenesis assay. (**h**) Quantification of autoradiography for labeled TAG was by ImageJ. *n* = 5, data are mean ± s.d. \**p* < 0.05.

To explore the molecular bases for altered lipolytic response of cool-adapted adipocytes, we performed immunoblot analyses and observed elevated expression of adipose triglyceride lipase (ATGL) and HSL in adipocytes cultured at 31°C (**Figure 6c**). Increased expression of lipolytic enzymes may seem paradoxical, given the observed reduction in basal lipolysis in cool-adapted cells; however, a lack of correlation between expression of ATGL or HSL and basal lipolysis has been observed previously (22). As expected, treatment of adipocytes cultured at 37°C with forskolin or CL increases phosphorylation of HSL at Ser563 and Ser660 (**Figure 6c**, **Figure S6a, S6b**). In cool-adapted adipocytes, we observed an excellent overall correlation between HSL phosphorylation and lipolysis - the increased phosphorylation of HSL at baseline is proportional to total HSL, and phosphorylation is substantially increased following treatment with forskolin but not CL.

Regulation of lipolysis is also dependent upon phosphorylation of perilipin by protein kinase A, which releases ABHD5 to co-activate ATGL (23). In this regard, analyses using a Phos-Tag gel system reveal that perilipin is dephosphorylated at baseline in adipocytes cultured at either temperature. However, whereas perilipin phosphorylation is increased by both forskolin and CL treatment in adipocytes cultured at 37°C, elevated phosphorylation is only observed with forskolin in adipocytes exposed to 31°C (**Figure 6c**, **Figure S6a, S6b**). One reason for impaired responsivity of cool-adapted adipocytes to CL is reduced expression of β3-AR, which is rapidly suppressed by exposure to 31°C (**Figure 3a**, **6d**). In addition, β2-adrenergic receptor expression is also rapidly suppressed (**Figure 6d**), suggesting a general resistance to adrenergic stimuli in cool-adapted cells. Taken together, exposure of adipocytes to 31°C blunts β3-adrenergic receptor expression and β3-AR-dependent lipolysis, but adaptation does not limit total lipolytic capacity.

### Adaptation to cool temperatures increases *de novo* lipogenesis and TAG synthesis

We next explored effects of cool adaptation on anabolic aspects of lipid metabolism. In addition to SCD1, exposure to 31°C up-regulates expression of other lipogenic proteins, such as fatty acid synthase (FASN) and acetyl-CoA carboxylase (ACC) (**Figure 6e**). Of note, *Fasn* mRNA expression is initially decreased after 1 day at 31°C, but is subsequently increased at 12 days of exposure (**Figure 3, 6f**). To evaluate functional effects of increased lipogenic gene expression with cool exposure, we incubated adipocytes cultured at 37°C or 31°C with [^14^C]-acetate for 12 hours (**Figure 6g, 6h**). Rates of *de novo* lipogenesis increased over eight days of cool adaptation, with increased radiolabel incorporation into TAG fractions of cellular lipid. Consistent with increased TAG synthesis in adipocytes adapted to 31°C, expression of *Dgat1*, which catalyzes the terminal step of TAG synthesis is also elevated (**Figure 6f** and **Supplemental data**). Taken together, these results indicate that cool adaptation up-regulates lipogenic gene and protein expression, and functionally increases *de novo* lipogenesis capacity and rates of TAG synthesis. Many *de novo* lipogenesis genes are regulated transcriptionally in adipocytes by sterol regulatory element-binding protein (SREBP-1c) and carbohydrate response element-binding protein (ChREBP) (24). Using loss-of-function approaches, we found that increased proteins involved in *de novo* lipogenesis in response to cool adaptation are regulated independently of these well-known transcription factors (**Figure S6c, S6d**).

### SCD1 activity is required for adipocytes to metabolically adapt to 31°C

Our studies have revealed that SCD1 mRNA and protein levels are up-regulated in cool-adapted adipocytes both in culture and *in vivo* (**Figures 1, 2,** and **6**). Further, we have shown that palmitic acid (C16:0), a substrate for SCD1, and oleic acid (C18:1), a product of this enzyme, are oxidized similarly in cool-adapted adipocytes (**Figure 5**). Thus, we next investigated the specific functional role of SCD1 in cool adaptation of adipocytes. Since cultured SCD1 knockout cells are prone to beiging (25) and have impaired lipid accumulation during adipogenesis (data not shown), we used a pharmacological approach for our studies. For adipocytes cultured at 37°C, inhibition of SCD1 activity for two days prior to the assay (**Figure 7a**) had no effect on OCR during Seahorse analyses (**Figure 7b-7f** and **S7a**). However, for adipocytes adapted to 31°C, SCD1 inhibition increased basal respiration, ATP-linked respiration, and proton leak, and reduced maximum respiration. Thus, spare respiratory capacity, which is defined as maximal OCR minus basal OCR in cool-adapted adipocytes, is specifically reduced in adipocytes cultured at 31°C (**Figure 7b-7f**). These data demonstrate that increased SCD1 activity is required for adipocytes to metabolically adapt to cooler temperatures.

**Figure 7.**
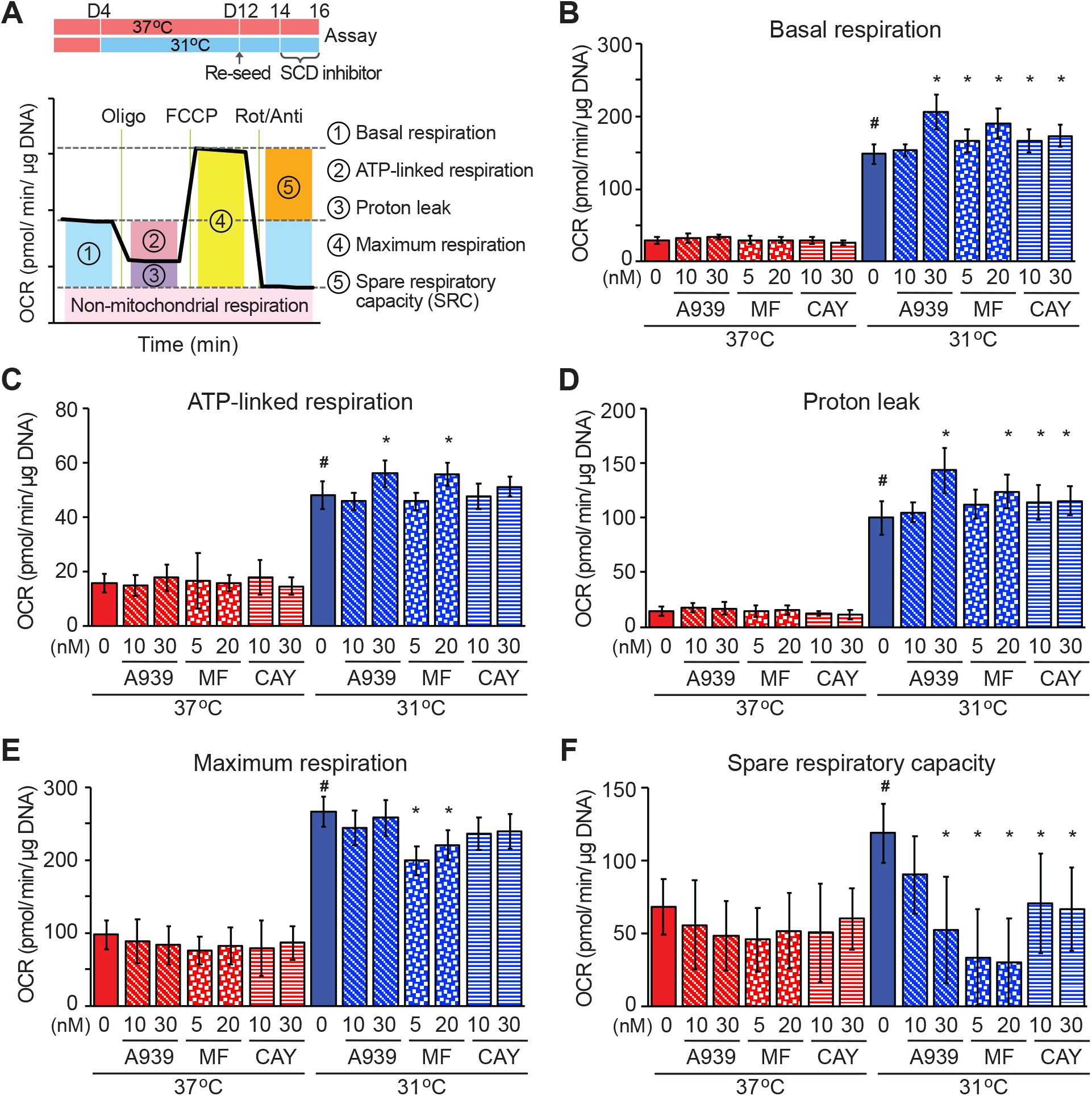
SCD1 activity is required for adipocytes to increase maximal OCR and spare respiratory capacity during adaptation to cool temperature. Differentiated adipocytes cultured at either 37°C or 31°C for 8 days were trypsinized and adipocytes were isolated by centrifugation. Adipocytes were seeded on Cell-Tak coated Seahorse XF96 microplates, then cultured overnight upside down at either 37°C or 31°C. Adipocytes were cultured with SCD inhibitors: A-939572, MF-438, or CAY10566 at the indicated temperature for two days before assay (*n* = 8-16 per group). (**a**) Summary of metabolic variables estimated. (**b**) Basal respiration. (**c**) ATP-linked respiration. (**d**) Proton leak. (**e**) Maximal respiration. (**f**) Spare respiratory capacity. Values are mean ± s.d. * Significantly different from vehicle control. *#* indicates a difference from 37°C vehicle control. Data shown is representative of at least 3 independent experiments.

## DISCUSSION

### Elevated oxygen consumption with cool adaptation is partially fueled by newly synthesized and stored fatty acids

Adipocytes exposed to 31°C are characterized by elevated oxygen consumption, with increased anabolic and catabolic lipid metabolic processes working in parallel. Although flux of glucose through glycolysis and the pentose phosphate shunt is suppressed, use of other nutrients to fuel oxidative metabolism, including pyruvate, glutamine and fatty acids is increased. Interestingly, cool adaptation of white adipocytes up-regulates *de novo* lipogenesis and expression of SCD1; thus, newly synthesized fatty acids are more highly desaturated prior to incorporation into TAG. Based on inhibition with etomoxir and atglistatin, NEFA from endogenous stores are the primary energy source for elevated oxygen consumption in adipocytes at 31°C. Cool-adapted adipocytes have higher demands for fatty acids hydrolyzed from TAG; however, they are also characterized by reduced secretion of glycerol and NEFA. Thus, cool adaptation of adipocytes is characterized by increased *de novo* lipogenesis, perhaps with pyruvate and amino acids as a source of carbon. TAG is then subject to partial hydrolysis in cool-adapted adipocytes, and whereas the newly hydrolyzed fatty acids are shuttled to mitochondria for oxidation, the acylglycerols are re-esterified to form TAG.

### Elevated SCD1 expression and monounsaturation of TAG lipids are cell-autonomous and required for elevated oxidative metabolism with cool adaptation

Previous studies in bacteria, plants, and poikilothermic animals have reported that cool temperatures increase lipid desaturation, whereas warm temperatures increase proportions of saturated lipids (26–28). This observation is largely considered to be due to homeoviscous adaptation, in which regulation of plasma membrane phospholipid desaturation is required to maintain appropriate viscosity and membrane function at various temperatures (29). However, our study reveals that compared to rats raised at thermoneutrality (29°C), distal BMAT of rats housed at 22°C exhibits increased TAG lipid unsaturation, whereas phospholipid composition is unaltered. A similar observation is found in subcutaneous WAT of shaved mice. A similarly disproportionate increase in lipid monounsaturation in TAG of cultured adipocytes was observed at 31°C, suggesting that elevated SCD1 preferentially influences composition of fatty acids incorporated into TAG versus phospholipids. Importantly, we demonstrate that SCD1 activity is required for maximal respiration and spare reserve capacity of adipocytes adapted to 31°C. These findings extend historical literature, which revealed a temperature gradient across the backfat of sows, and revealed that soft lard from the outer, cooler layer of subcutaneous fat contains increased unsaturated lipids, particularly oleate (14). Together, these findings suggest an evolutionarily-conserved role for lipid desaturation of TAG in metabolic adaptation to cool cellular environments.

Oleic acid and palmitoleic acid have previously been reported to induce expression of OXPHOS subunits 3T3-L1 adipocytes and fatty acid oxidation in skeletal muscle cells (30, 31). One intriguing possibility is that the observed increases in OXPHOS subunits and β-oxidation are regulated by increased production of monounsaturated lipids with cool adaptation. Indeed, we observed a higher proportion of the SCD1 product oleic acid (C18:1, n-9) esterified in TAG of distal tibia and CV BMAT, which exist at temperatures lower than those found at the body core. Instead, it is the SCD1 product palmitoleic acid (C16:1, n-7) that is dramatically increased by cool exposure of cultured adipocytes. Coincident with improved mitochondrial function, we observed increased expression of OXPHOS subunits with cool adaptation. In our studies, complexes I, II, and III were up-regulated in adipocytes at 31°C, whereas previous work in differentiated 3T3-L1 cells reported that palmitoleic acid-induced oxygen consumption and complexes II, III, and V (31).

### Identification of adaptive thermal responses specific to white adipocytes

Previous studies using clonal, immortalized cell lines and bone marrow-derived MSCs have suggested that in adipocytes, cool temperatures can activate the expression of thermogenic genes, such as *Ucp1* and *Pgc1a*, in a cell-autonomous manner (7, 32). In this work, we have demonstrated that white adipocytes isolated from the epididymal depot, and adipocytes differentiated from MSCs or SVCs from WAT, do not up-regulate expression of UCP1 in response to cool temperatures. Moreover, our RNA-seq dataset did not reveal induction of canonical browning/beiging genes in these cells. It is possible that the strategies used by specific adipocyte models to adapt to cool temperatures depend on the propensity of individual cell types for browning/beiging.

From an energy utilization viewpoint, cool-adapted adipocytes possess high fatty acid oxidation capacity and preferentially use fatty acids over glucose as a fuel source. This metabolic shift for white adipocytes is distinct from other types of adipocytes; brown/beige adipocytes are characterized by high glucose uptake and oxidative capacity in “glycolytic beige fat” (33), and both glucose and fatty acids are used as substrates in conventional thermogenic brown and beige adipocytes following cold stimulation (34–37). Importantly, exposure to cool temperatures does not influence the expression of genes that define the adipocyte phenotype, nor are there differences observed in adipocyte morphology, including the appearance of brown or beige characteristics. Thus, our results demonstrate that white adipocytes adapted to cool temperatures have unique molecular and metabolic characteristics that distinguish them from counterparts cultured at 37°C and from brown/beige adipocytes. Our study raises the possibility that cool environmental temperature may be an unappreciated factor in stimulating the distinct cellular and physiological features of adipocytes found within different WAT depots.

## METHODS

### Animals

Sprague Dawley rats were obtained from Envigo and 16 females were bred. Two days after giving birth, eight dams and their litters remained in thermal chambers at 22°C and eight were transferred to thermal chambers at 29°C (i.e. thermoneutrality). Eleven weeks later, male and female rats were sacrificed. C57BL/6J mice (JAX Labs) were bred in thermal chambers at 22°C or 29°C. Male mice were housed from birth to 13 weeks either at 22°C or at 29°C. After weaning, posterior mouse hair was removed using Nair. Mice were single-housed with minimal bedding and no nesting materials. Subdermal temperature was measured of mice housed at 22°C or 29°C using a rodent thermometer BIO-TK8851 (Bioseb Lab Instruments) under anesthesia. Temperature measurement was finished within 5 min of anesthesia to avoid temperature-lowing effects. All animal studies were performed in compliance with policies of the University of Michigan Institutional Animal Care and Use Committee.

### Cell Culture

Primary mesenchymal stem cells (MSCs) were isolated from the ears of wildtype C57BL/6J, UCP1-KO (Ucp1^tm1Kz^/J) (The Jackson Laboratory), and ChREBP KO (kindly gifted by Dr. Lei Yin at the University of Michigan) mice as previously described (38). At two days post-confluence, adipogenesis was induced with 0.5 mM methylisobutylxanthine, 1 μM dexamethasone, 5 μg/ml insulin and 5 μM rosiglitazone (Cayman Chemical) in DMEM:F12 containing 10% FBS. From day 2 to 4 of differentiation, cells were fed with fresh DMEM:F12 medium containing 10% FBS, 5 μg/ml insulin and 5 μM rosiglitazone. Thereafter, cells were maintained in DMEM:F12 containing 10% FBS.

### Reagents

Reagents used were as follows; adenosine 5’-diphosphate (Sigma-Aldrich), A-939572 19123 (Cayman Chemical), CAY10566 (Cayman Chemical), Cell-Tak (Corning), CL-316,243 (Tocris), C75 (Cayman Chemical), etomoxir (Cayman Chemical), forskolin (Tocris), MF-438 (Millipore Sigma), octanoic acid (Sigma-Aldrich), oleic acid (Sigma-Aldrich), rotenone (Tocris), palmitoleic acid (Sigma-Aldrich), palmitoyl carnitine (Sigma-Aldrich), pyruvate (Sigma-Aldrich), sodium palmitate (Sigma-Aldrich), 5-(tetradecyloxy)-2-furoic acid (TOFA) (Cayman Chemical)

### Adipocyte and stromal-vascular cell fractionation

Gluteal WAT (GluWAT) and epididymal WAT (eWAT) were excised from mice, minced with scissors and digested for 1 h at 37°C in 2 mg/ml collagenase type I (Worthington Biochemical) in Krebs-Ringer-HEPES (KRH; pH 7.4) buffer containing 3% fatty acid-free bovine serum albumin (BSA; Gold Biotechnology, St. Louis, NJ), 1 g/L glucose, and 500 nM adenosine. The resulting cell suspensions were filtered through 100 μm cell strainers and centrifuged at 100 x g for 8 min to separate the stromal-vascular fraction and the buoyant adipocytes. Floating adipocytes were washed and cultured with DMEM:F12 containing 10% FBS.

### Isolation and differentiation of adipocyte precursors

Mesenchymal precursors were isolated from ears of mice of the indicated genotypes, as previously described (18). Cells were maintained in 5% CO_2_ and DMEM/F12 1:1 media (Gibco; Invitrogen) supplemented with 15% FBS (Atlas Biologicals), primocin (InVivoGen), and 10 ng/ml recombinant bFGF (PeproTech). For induction of adipogenesis, recombinant bFGF was removed and replaced with 10% FBS containing 0.5 mM methylisobutylxanthine, 1 μM dexamethasone, 5 μg/ml insulin, and 5 μM troglitazone. On day 2, cells were fed 5 μg/ml insulin plus 5 μM troglitazone. On day 4 and every 2 days thereafter, cells were fed with 10% FBS. At day 4 of differentiation, plates of adipocytes were either kept at 37°C, or moved to another incubator maintained at 31°C. Of note, since adipocyte culture media contains both sodium-bicarbonate and HEFPES, pH change are unlikely to contribute to temperature-induced effects on adipocyte gene expression.

### Lipolysis

Cultured adipocytes were washed 1x with PBS and then incubated in Hank’s Balanced Salt Solution (HBSS; Thermo Fisher Scientific) containing 2% BSA for 1h at 37°C. Cells were treated with forskolin or CL-316,243 for 6 h. Secretion of glycerol and non-esterified fatty acid from cultured adipocytes into HBSS was determined with assay kits from Sigma-Aldrich (FG0100) and FUJIFILM Wako Diagnostics-NEFA Reagent (NEFA-HR(2)), respectively.

### *De novo* lipogenesis assay

Cultured adipocytes were incubated overnight in fresh serum-free DMEM:F12 medium prior to measurement of *de novo* lipogenesis. Cells were then incubated in fresh DMEM:F12 medium (containing 0.5 mM sodium pyruvate, 0.5 mM L-glutamine, 2.5 mM glucose) supplemented with 1% fatty acid-free BSA, and containing 0.5 μCi [^14^C]-acetate (PerkinElmer) and 5 μM sodium acetate for 12 h at either 37°C or 31°C. At the end of the indicated incubation time, adipocytes were harvested, and lipids extracted for analyses by thin layer chromatography.

### β-oxidation assay

Cultured adipocytes were washed 1x with PBS and then incubated with DMEM (following manufacturer’s guidelines; catalog #D5030; Sigma-Aldrich) containing BSA-conjugated ~3 μCi/ml of either [9,10-3H(N)]-palmitic acid, [9,10-3H(N)]-oleic Acid (PerkinElmer) or n-[2,2’,3,3’-3H] octanoic acid (American Radiolabeling Chemicals) and 22 μM of unlabeled non-esterified fatty acids respectively (Sigma-Aldrich). For long-chain fatty acid oxidation, assay media was contained with 200 μM L-carnitine (Sigma-Aldrich), and a subset of wells were treated with 40 μM etomoxir (Cayman Chemical) 2 h before and duration of the assay. After 3 h, conditioned media from cells was passed through columns containing AG1-X8 Anion Exchange Resin (Bio-Rad) and collected in scintillation vials, then mixed with scintillation cocktail Bio-Safe II (RPI Research Product International). Radioactivity in the supernatant was measured using a scintillation counter.

### Extracellular flux assay

Cellular and mitochondrial OCR were determined using the Seahorse XF96 Extracellular Flux Analyzer (Agilent). To measure OCR on cool adapted adipocytes, we used two different approaches, and which gave similar results. In the first approach, a 96 well cell culture microplate for Seahorse was cut into half, and then ~480 MSCs were seeded into each well. Four days after seeding, adipogenesis was induced at 37°C same as described above. Four days after induction of differentiation, adipocytes in half of the plate were cultured at 31°C whereas the other half were incubated at 37°C for 12 days. At this point, the plate was rejoined in order to evaluate oxygen consumption at 31°C or at 37°C. In the second approach, differentiated adipocytes cultured at either 37°C or 31°C for 8 days (four days before the assay) were trypsinized and isolated by centrifugation. Adipocytes were seeded on Cell-Tak coated Seahorse XF96 microplates, then cultured overnight upside down at either 37°C or 31°C. One of the advantages of the first approach is that adipocytes are cultured on the same plates for the entire experiment. However, it is difficult in long term culture to control cell numbers and thus basal OCR. Although we favor use of the second approach, and in our hands the replated adipocytes appear to maintain their differentiate morphology, one concern is the potential for adipocyte dedifferentiation. On the day of assay, the cells were washed and incubated with unbuffered XF assay medium (Agilent;103575) contained 5 mM glucose, 0.2 mM pyruvate and 1 mM glutamine and placed the cell culture microplate into either 31°C or 37°C non-CO2 incubator for 1.5 h prior to the assay. The sensor cartridge was loaded with oligomycin (Port A), FCCP (Port B), and rotenone/antimycin A (Port C) (XF Cell Mito Stress Test Kit, Agilent) to achieve final concentrations of 1 μM, 1 μM, and 0.5 μM, respectively. Other procedures for the assay were performed by following the manufacturer’s instructions. OCR was measured and normalized to DNA. Isolated mitochondria (5 μg protein) from MSC-derived adipocytes were seeded to XF96 microplates, and OCR was assayed as described above. Final concentrations of reagents are following; 1mM Adenosine 5’-diphosphate (ADP), 10 mM succinate, 2 μM rotenone, 10 mM pyruvate, 1 mM malate, 40 μM palmitoylcarnitine, For the parametric equations: Basal respiration = (initial respiration) - (non-mitochondrial respiration); ATP-linked respiration = (last OCR before oligomycin) - (minimum OCR after oligomycin); Proton leak = (minimum OCR after oligomycin) - (non-mitochondrial OCR); Maximal respiration = (maximum rate after FCCP injection) - (non-mitochondrial respiration); SRC = (maximal respiration) - (basal respiration).

### Isolation of mitochondria from adipose tissue and cultured MSC adipocytes

Isolation of mitochondria was as described previously (18). Briefly, cultured MSC-derived adipocytes or adipocytes isolated from gluteal WAT were homogenized using a Potter-Elvehjem homogenizer and centrifuged at 800 g for 10 min at 4°C. The supernatant was then centrifuged for 15 min at 8,000 g at 4°C, and the pellet was washed with ice-cold buffer. After centrifugation at 7,000 g for 10 min at 4°C, the pellet containing mitochondria was resuspended for analyses.

### Metabolomics

Adipocytes from MSC were pre-incubated with fresh DMEM (#D5030; Sigma-Aldrich) with unlabeled glucose (5 mM), pyruvate (0.2 mM) and glutamine (1 mM) for 2 h. For incubation in tracer-labeled media, a media change was performed. Substrate concentrations were kept constant except for the substitution of unlabeled metabolites with either ^13^C_6_ glucose (Sigma-Aldrich), 2,3-^13^C_2_ pyruvate (Cambridge Isotope) or ^13^C_5_ glutamate (Sigma-Aldrich) for 1 hour. Cells were then rapidly (<5 seconds) rinsed with 150 mM ammonium acetate, and snap-frozen by addition of liquid nitrogen directly to the cell plate. Metabolites from frozen cells were extracted by adding ice-cold 8:1:1 HPLC grade methanol:chloroform:water to the frozen cell plate, scraping to detach and lyse cells, transferring the suspension to microcentrifuge tubes, and centrifuging for 5 min at 15k RCF to remove pellet debris. Polar metabolites were analyzed in the supernatant by hydrophilic interaction chromatography-electrospray time of flight mass spectrometry (HILIC-ESI-TOF) as described previously (39).

### Lipid extraction

Cultured adipocytes were washed twice with PBS and then suspended in 500 μl of a 1:2.5 methanol/water mixture. The cell suspension was transferred to a borosilicate glass tube. Wells were rinsed with 500 μl of the 1:2.5 methanol/water mixture and the volume was transferred to a glass tube, then vortexed after adding 375 μl chloroform. Then another 375 μl chloroform and 375 μl 0.9% NaCl were added to each tube, which was then vortexed vigorously and centrifuged at 2,500 rpm for 20 min at 4°C. The lower organic chloroform layer containing total lipids was transferred to a new tube and stored at −20°C until analyzed. For the BMAT and WAT, tissue was transferred to a borosilicate glass tube after crushing on dry ice. Add 1000 μl of the 1:2.5 methanol/water mixture 375 μl chloroform, then homogenize with immersion tissue grinder. Then another 375 μl chloroform and 375 μl 0.9% NaCl were added to each tube, which was then vortexed vigorously and centrifuged at 2,500 rpm for 20 min at 4°C. The next step is the same as the process as described above for the cells.

### Chromatographic separation of TAG and phospholipids

TAG and phospholipids were separated from the total lipid extract of cultured adipocytes and adipose tissue, by thin-layer chromatography applying the lipids as a band on a 20 x 20 cm silica gel thin-layer chromatography plate (silica gel 60, Millipore Sigma). The plate was developed with hexane-diethyl ether-acetic acid (80-20-1.5, v/v). The TAG and phospholipids were identified with respect to the retention flow (rf) of the authentic TAG and phospholipid standard applied on the same plate. TAG and phospholipid bands were scraped from the thin-layer chromatography plate, extracted with chloroform and the lipids were subjected to derivatization as follows.

### Trans-esterification with BF_3_-methanol and gas chromatography

Fatty acids in the extracted lipids were derivatized into their methyl esters by trans-esterification with boron trifluoride-methanol. The derivatized methyl esters were re-dissolved in a small volume of hexane and purified by thin-layer chromatography using n-hexane-diethyl ether-acetic acid (50:50:2, v/v/v) as the developing solvents. After development, plates were dried and sprayed with Premulin. Fatty acid methyl ester bands were identified under ultraviolet light by comparing the retention flow of methyl heptadecanoate (C17:0) standard (retention flow, 0.67) applied side-by-side on the same plate. Methyl esters were extracted from thin-layer chromatography powder with diethyl ether concentrated under nitrogen and re-dissolved in 100 μl hexane. The lipids’ fatty acid compositions were analyzed by gas chromatography (GC) as follows. FAMEs were analyzed with 1 μl sample injection on an Agilent GC machine, model 6890N equipped with a flame ionization detector, an autosampler model 7693 and a ChemStation software for data analysis. An Agilent HP 88 with a 30 m GC column with a 0.25 mm inner diameter and 0.20 mm film thickness was used. Hydrogen was used as a carrier gas and nitrogen was used as a makeup gas. The analyses were conducted with a temperature programming of 125-220°C. The fatty acid components within unknown samples were identified with respect to the retention times of authentic standard methyl ester mixtures run side by side. The fatty acid components were quantified with respect to the known amount of internal standard added and the calibration ratio was derived from each fatty acid of a standard methyl ester mixture and methyl heptadecanoate internal standard. The coefficient of variation for GC analyses was within 2.3-3.7%.

### qPCR to quantify mtDNA

DNA was prepared from cells using Gentra Puregene Kits including RNase A treatment (Qiagen). mtDNA copy number per nuclear genome in MSC-derived adipocytes was quantified as described previously (18). Primer sequences for PCR are in Supplemental Table 1.

### mRNA quantification by RT-PCR

RNA isolation, reverse transcription and quantitative PCR were performed as previously described (18). Primer sequences for real-time RT-PCR are in Supplemental Table 1.

### RNA-Sequencing

Total RNA was isolated and purified as described above. After DNase treatment, samples were submitted to the University of Michigan Advanced Genomics Core for quality control, library preparation, and sequencing using the Illumina Hi-Seq platform. Read files were downloaded and concentrated into a single fastq file for each sample. Quality of raw read data was checked using FastQC (version v0.11.30) to identify features of the data that may indicate quality problems (i.e. low-quality scores, over-represented sequences, inappropriate GC content). The Tuxedo Suite software package was used for alignment, differential expression analysis, and post-analysis diagnostics. Briefly, reads were aligned to the UCSC reference genome using TopHat (version 2.0.13) and Bowtie2 (version 2.2.1). FastQC was used for the second round of quality control post-alignment to ensure that only high-quality data was input to expression quantitation and differential expression analysis. Differential expression analysis was done using two distinct methods to check for analysis consistency: Cufflinks/CuffDiff and HTSeq/DESeq2, using UCSC build mm10 as the reference genome sequence. Plots were generated using variations or alternative representations of native DESeq2 plotting functions, ggplot2, plotly, and other packages within the R environment.

### Pathway Analysis

Pathway analysis was conducted on ranked lists of log2 fold change using Gene Set Enrichment Analysis (GSEA) v4.0.3 by the Broad Institute. Prior to analyses, mouse gene symbols were remapped to human ortholog symbols using chip annotation files. The mouse genes that did not have equivalent human orthologs were excluded from the analysis. The output from DESeq2 analysis was used to generate a list of genes ranked by the metric −log10 FDR * log2 fold change. The resulting list was run through pre-ranked GSEA using the Molecular Signatures Database v7.1 (H, hallmark gene sets). Enriched pathways were defined by an FDR < 0.05. The normalized enrichment score (NES) is the primary metric from GSEA for evaluating the magnitude of differentially expressed pathways. Pathway impact analysis was conducted on AdvaitaBio’s iPathwayGuide to identify enriched KEGG pathways. The ggplot2 package in R was used for further visualization of the enriched pathways.

### Immunoblot

After lysis in 1 % NP-40, 120 mM NaCl, 50 mM Tris-HCl; pH 7.4, 50 mM NaF, 2 mM EDTA, 1x protease inhibitor cocktail (Sigma-Aldrich), protein concentrations of lysates after centrifugation were measured by BCA protein assay (Thermo Fisher Scientific). Lysates were diluted to equal protein concentrations in lysis buffer and then boiled in SDS sample buffer (20 mM Tris; pH 6.8, 2% SDS, 0.01% bromophenol blue, 10% glycerol, 5% 2-mercaptoethanol) and subjected to SDS-PAGE and immunoblotting according to standard techniques. Separation of phosphorylated proteins using the Phos-Tag gel (FUJIFILM Wako Chemicals) was according to manufacturer’s instructions. Quantification of protein expression was done using ImageJ software. Antibodies used were as follows; ACC (Cell Signaling Technology #3662), p-ACC(Ser79) (Cell Signaling Technology #3661), Adiponectin (Sigma Aldrich A6354), ACAA2 (MCKAT) (Thermo Fisher Scientific A21984), ATGL (Cell Signaling Technology #2138), β-actin (Cell Signaling Technology #4970), CPT1A (Abcam ab128568), CHREBP (Novus Biologicals NB400-135), ERK2 (Santa Cruz Biotechnology sc-1647), FASN (Abcam ab22759), FABP4 (R&D #1443), HSL (Cell Signaling Technology #4107), p-HSL (Ser563) (Cell Signaling Technology #4139), p-HSL (Ser660) (Cell Signaling Technology #4126). HADHB (MTPβ) (Novus Biologicals NBP1-54750), HSP70 (BD Transduction lab 610607), HSP90 (BD Transduction lab 610418), Laminin (Novus Biologicals, NB300-144), OXPHOS Cocktail (Abcam ab110413), Perilipin (Abcam Ab3526), PMP70 (Thermo Fisher Scientific PA1650), PEX5 (Thermo Fisher Scientific PA558716), PPARg (Millipore MAB3872), SCD1 (Cell Signaling Technology #2438), SREBP1 (ThermoFisher Invitrogen 2A4), Tubulin (Invitrogen MA1-80017), UCP1 (Alpha Diagnostic UCP11-A), VDAC1 (Abcam ab15895).

### Statistics

All data are presented as mean ± S.D. When comparing two groups, significance was determined using two-tailed Student’s t-test. When comparing multiple experimental groups, an analysis of variance (ANOVA) was followed by post hoc analyses with Dunnett’s or Sidak’s test, as appropriate. Differences were considered significant at *p* < 0.05 and are indicated with asterisks. For metabolomics data analyses, the proportion of labeling at each carbon position was calculated by dividing each species by total sum of peak areas of all labeled positions. The proportion of data is well-known to follow a beta distribution. Beta regression model is an extension of the Generalized Linear Model with an assumption that the response variable follows a beta distribution with values in standard unit interval (0,1) (40). Our flux proportion data contains zero and one values depending on the labeling. Therefore, our response variable does not fall under the standard unit interval (0,1). Ospina and Ferrari proposed a more general class of zero-or-one inflated beta regression models for continuous proportions (41). We proposed a zero-and-one inflated model, which was applied using R package GAMLSS (42). To apply multiple testing correction, the significance is measured by a q-value calculated using the algorithm developed by Storey and Tibshirani (43). All data analysis was performed in R.

## ACKNOWLEDGEMENTS

This work was supported by grants from the NIH to OAM (RO1 DK62876; R24 DK092759; R01DK126230), DPB (T32 HD007505; T32 GM007863), CAC (T32 DK101357), and SMR (T32 GM835326; F31 DK12272301, and from the American Diabetes Association (1-18-PDF-087) to ZL and CAC (1-18-PDF-064). This work was also supported by Agilent. We are grateful for the support of several core facilities, including the University of Michigan Advanced Genomics Core, Michigan Regional Comprehensive Metabolomics Resource Core (U24 DK097153), and Adipose Tissue Core of the MNORC (P30 DK089503). We thank Soni Tanu for statistical analysis of lipidomics and metabolomics data, Catherine Salamon, Jihan Khandaker, Johena Sanyal, Ahmad Mustafa, Heidi Baum, Angela Wiggins, Becky Tagett, Kathryn Tormos, George Rogers, Genevieve Van de Bittner, and Xie Yue for technical assistance, and Takeshi Akama and members of the MacDougald lab for helpful discussions and assistance.

## AUTHOR CONTRIBUTIONS

H.M., C.D., and O.A.M. conceived the project, designed the experiments. H.M. and O.A.M wrote the manuscript. H.M. performed the majority of the experiments, data analyses, and manuscript preparation. C.D. performed lipid profiling. A.N. performed analyses of RNA-seq data. A.D. performed lipidomics assay. A.B., T.C., and S.P. assisted with performance of experiments. Z.L., S.R., C.C, D.B, J.H, and B.L. assisted with performance of large-scale animal studies. C.E. K.O. and K.I. contributed intellectually to experiments and provided key feedback.

## Supplemental Information

**Supplemental Figure 1.**
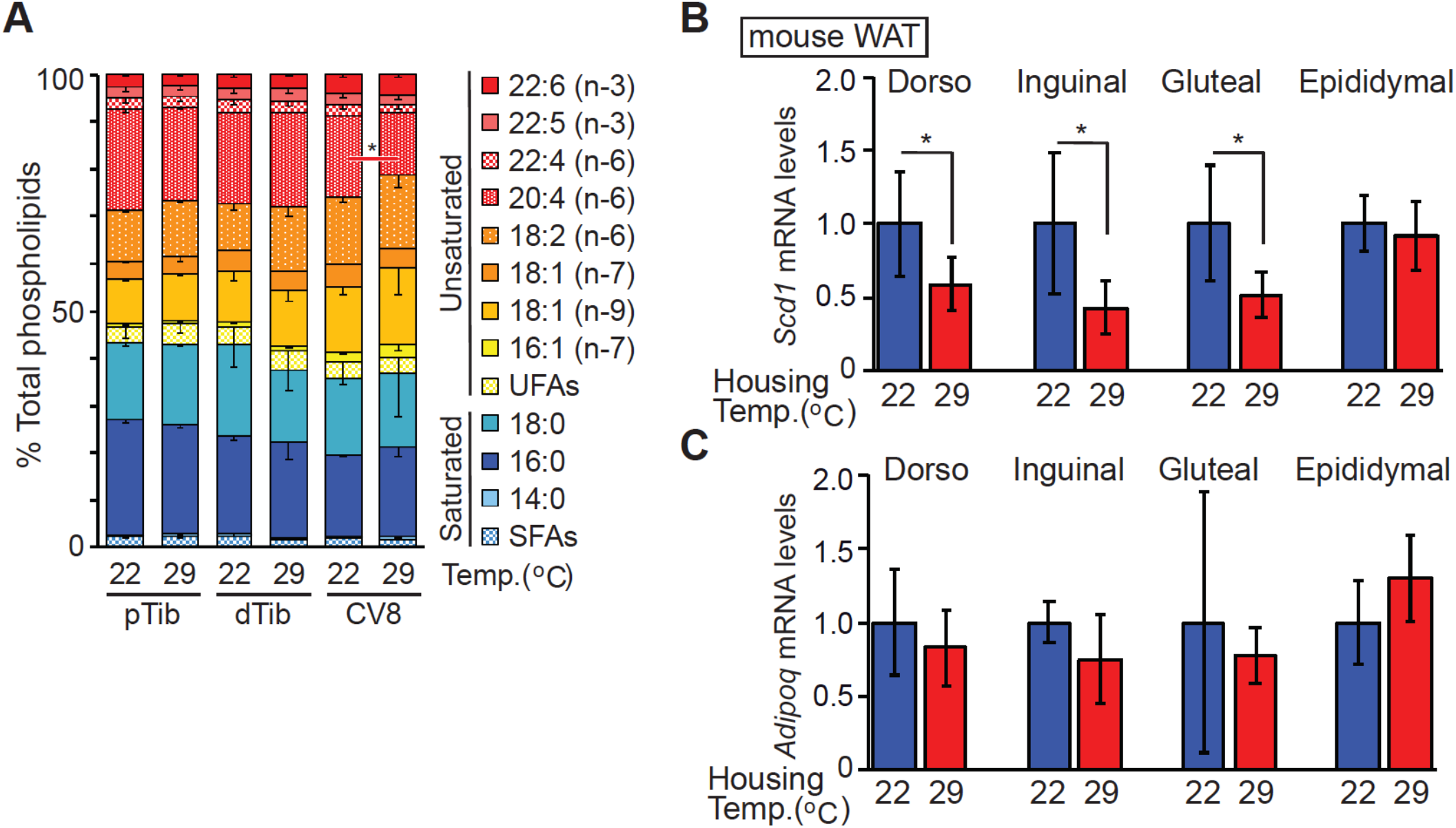
(**A**) Rats were housed from birth to 11 weeks of age at room temperature (22°C) or thermoneutrality (29°C). Lipid composition of phospholipids for proximal (pTib) and distal tibia (dTib), and caudal vertebra-8 (CV8) was determined by gas chromatography (n = 6). (**B** and **C**) Mice were housed from birth to 13 weeks either at 22°C or at 29°C without posterior hair after weaning. Whereas *Scd1* mRNA expression is elevated in subcutaneous WAT depots of mice at 22°C (**B**), *Adipoq* expression was not altered(**C**). Gene expression was normalized to geometric mean value of *Hprt*, *Tbp*, *Gapdh*, *Rpl32* and *Ppia,* and was expressed relative to 37°C control (*n* = 8-9). For panels (**A**) - (**C**), values are mean ± s.d. \**p*<0.05. Data shown is representative of at least 3 independent experiments.

**Supplemental Figure 2.**
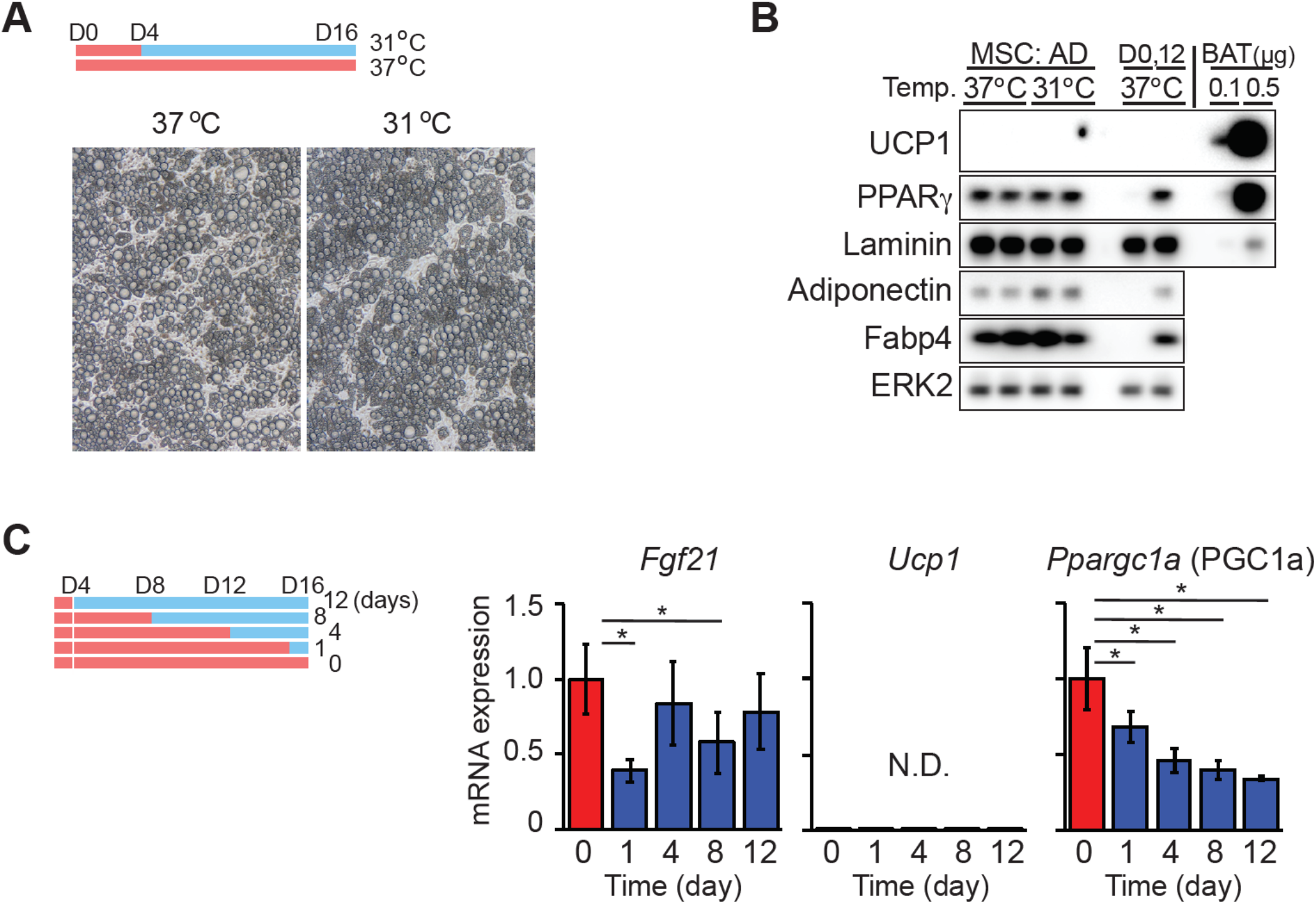
(**A**) Cultured MSC adipocytes adapt to 31°C without morphological changes. Four days after differentiation, adipocytes were moved to the indicated temperatures for 12 days. (**B**) Adipocyte markers are expressed similarly between control cells cultured at 37°C and those incubated at 37°C from day 4 to 16. For adipocyte samples, 11 μg of protein lysate analyzed per lane, whereas for brown adipose tissue, 0.1 or 0.5 μg lysate was included as a positive control for UCP1. (**C**). Beige adipocyte markers were not induced by exposure of cultured adipocytes to 31°C for the indicated days (n = 6). Gene expression was normalized to geometric mean value of *Hprt*, *Tbp*, *Ppia,* and was expressed relative to 37°C control (31°C for 0 day) (*n* = 4). * indicates significance at *p* < 0.05.

**Supplemental Figure 3.**
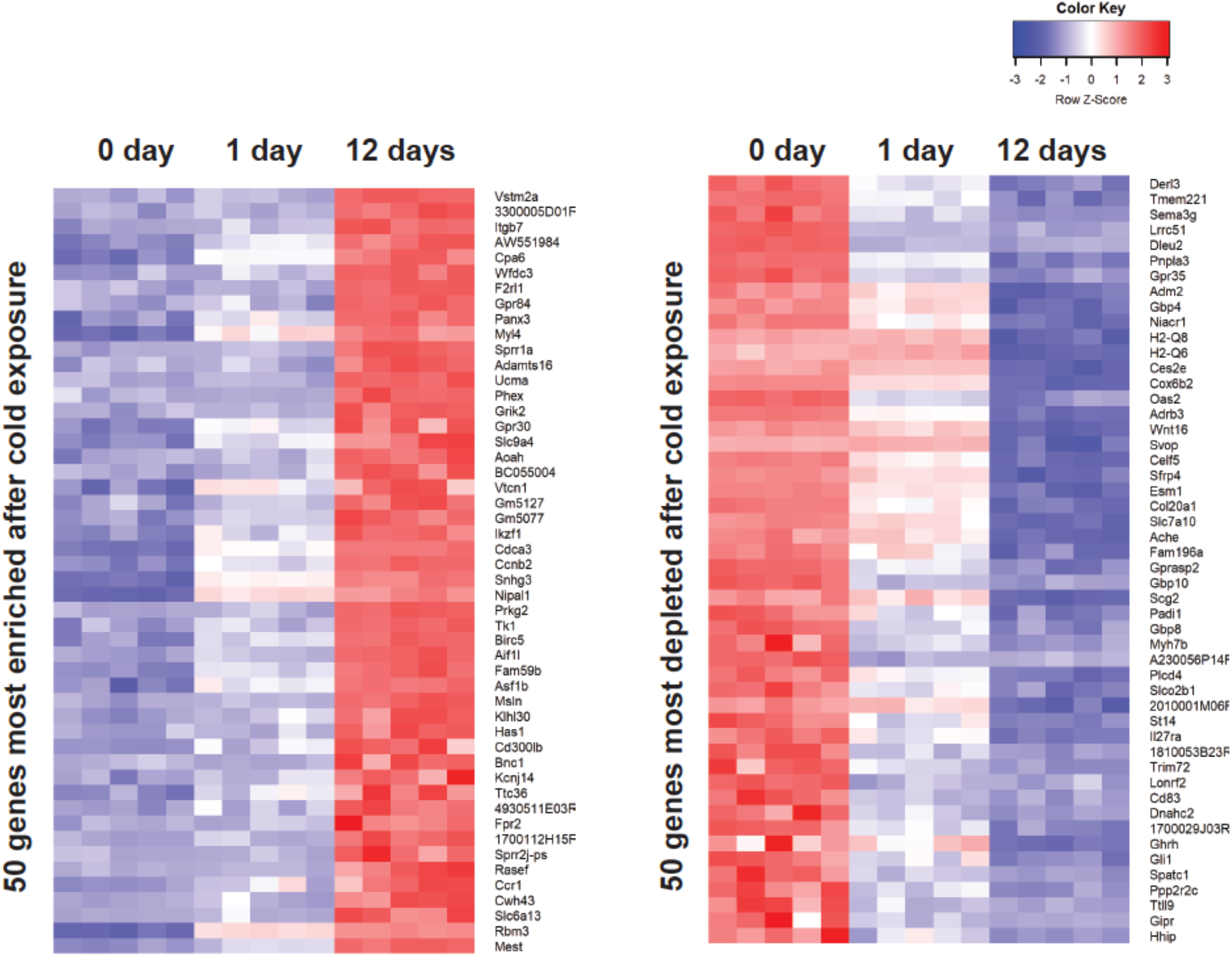
Heat map of top 50 enriched and top 50 depleted genes in 12-day cool exposed MSC adipocytes. Color key based on rlog-transformed read count values and significance was defined by an FDR < 0.01.

**Supplemental Figure 4.**
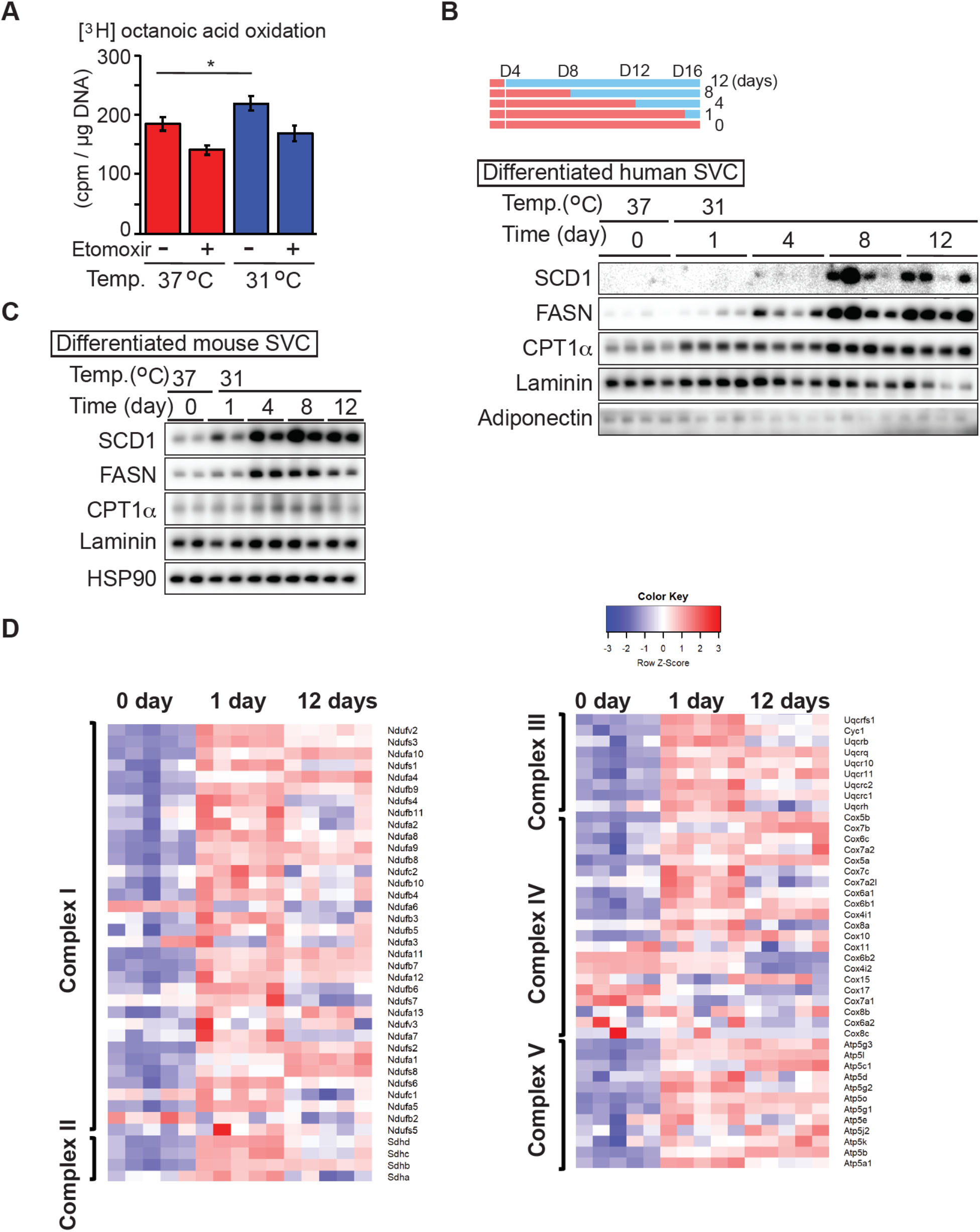
(**A**) Release of tritiated water from adipocytes treated with labeled octanoic acid. Adipocytes cultured at indicated temperature for 12 days were incubated with tritiated octanoic acid for 3 hrs in the presence and absence of etomoxir (*n* = 5). (**B** and **C**) Cool adaptation increases enzymes involved in synthesis and degradation of non-esterified fatty acids. Lysates were collected after the indicated days of cool adaptation. Stromal-vascular cells (SVC) of SVC from human (**B**) or eWAT from C57BL/6J mice **(C)** were differentiated into adipocytes. Human white preadipocytes (kindly provided by Dr. Shingo Kajimura; UCSF) were differentiated as previously described (44). (**D**) Oxidative phosphorylation genes are upregulated at the mRNA level. Heat map of genes involved in complexes I, II, III, VI, and V were constructed from KEGG map of oxidative phosphorylation genes (mmu00190).

**Supplemental Figure 5.**
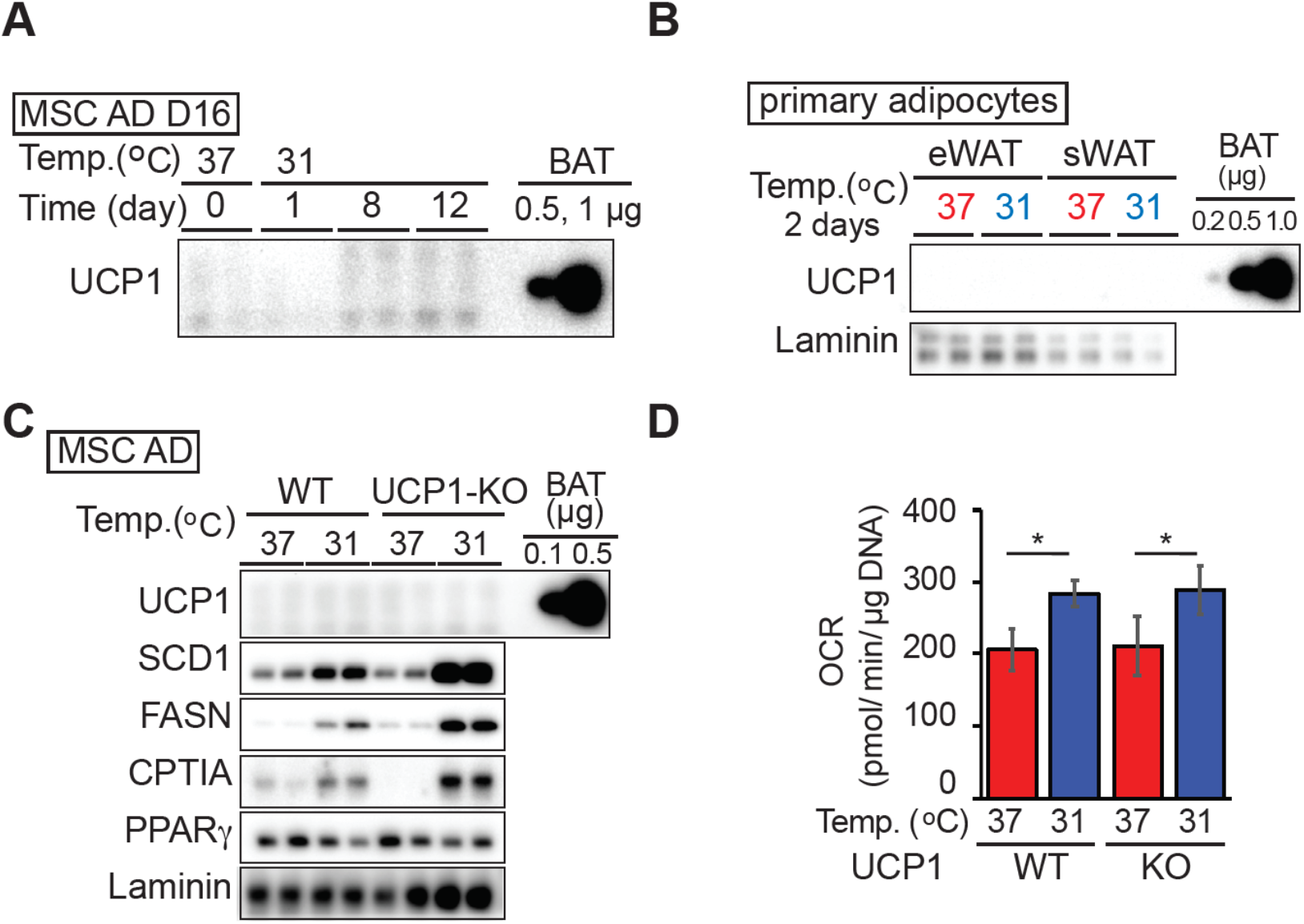
(**A**) Expression of UCP1 in adipocytes adapted to 31°C for the indicated days. 11 μg of adipocyte or 0.5 or 1 μg BAT lysate was evaluated by immunoblot for UCP1. (**B**) Primary adipocytes isolated from epididymal white adipose tissue (eWAT) or subcutaneous gluteal WAT (sWAT) by collagenase digestion were cultured floating at either 37°C or 31°C for 2 days. Brown adipose tissue lysate (0.2, 0.5 or 1 μg) was used as a positive control for UCP1. (**C**) Cool adaptation increases enzymes involved in synthesis and degradation of non-esterified fatty acids in adipocytes derived from UCP1 knockout mice. (**D**) Elevated basal OCR of adipocytes at 31°C is UCP1 independent. MSC adipocytes derived from wild type or UCP1 knockout mice were cultured at 31°C or 37°C for 12 days and basal OCR evaluated (*n* = 8).

**Supplemental Figure 6.**
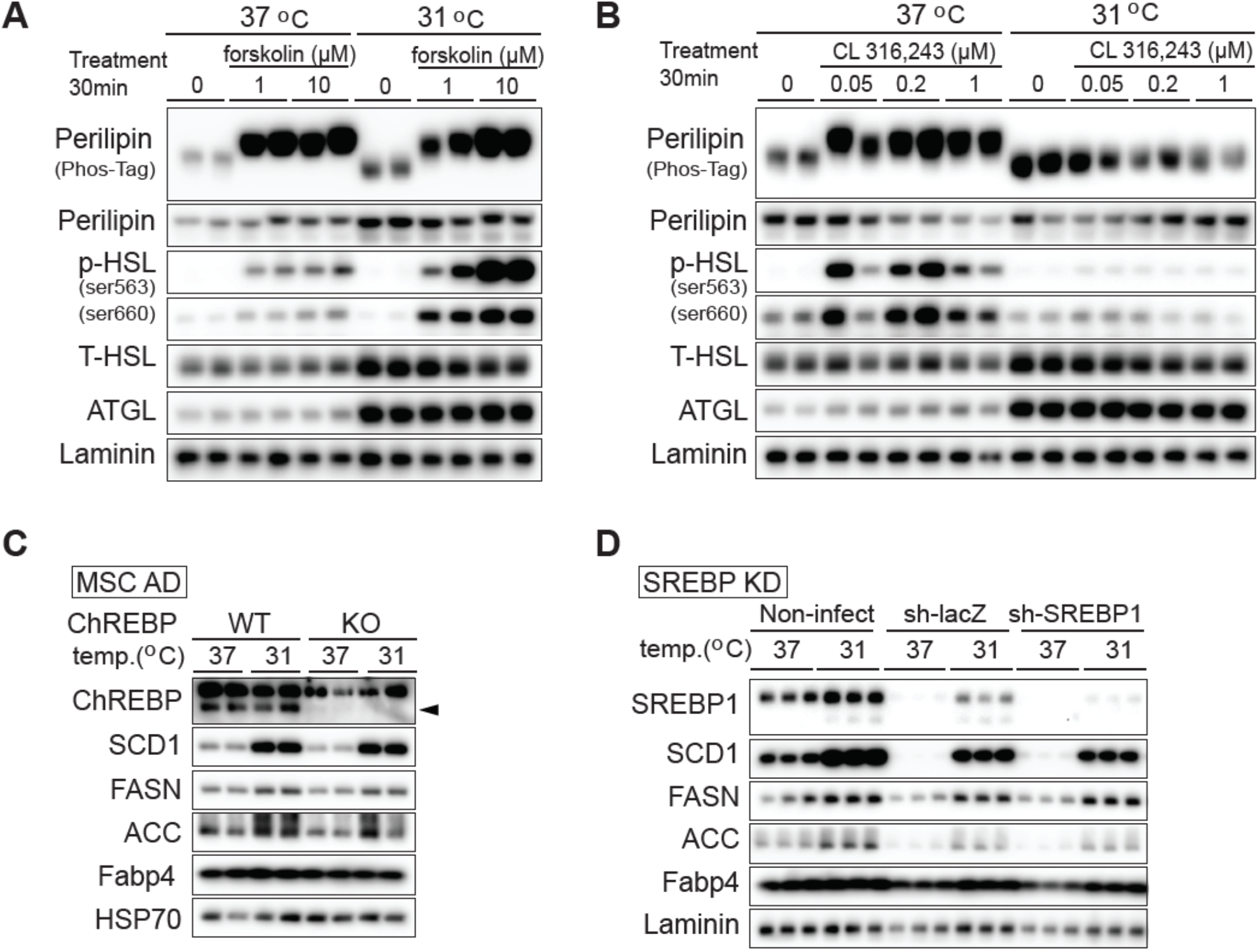
(**A** and **B**) Cool adaptation increases the expression of HSL and ATGL but decreases phosphorylation of perilipin and HSL in response to CL-316,243. Mature adipocytes were adapted to 31°C for 12 days or remained at 37°C. Adipocytes were treated with **(A)** vehicle, 1 μM or 10 μM forskolin, or **(B)** vehicle, 0.05 μM, 0.2 μM or 10 μM CL-316,243 for 30 min. Lysates were collected for immunoblot analyses. (**C**) Induction of enzymes involved in *de novo* lipogenesis with adaptation to 31°C is independent of ChREBP. Adipocytes derived from wildtype or ChREBP knockout mice were cultured at 31°C or 37°C for 4 days before collecting samples. Although ChREBP-deficiency impairs adipocytes differentiation (45), addition of rosiglitazone to differentiation media resulted in robust adipogenesis of both sets of cells, as has been reported (46). (**D**) At day six of MSC differentiation, adipocytes were infected in serum-free medium with adeno-sh*LacZ* control or adeno-sh*Sreb1* to induce gene knockdown. Adipocytes were then allowed to recover following infection and were cultured at the indicated temperature from day 8 to day 12.

**Supplemental Figure 7.**
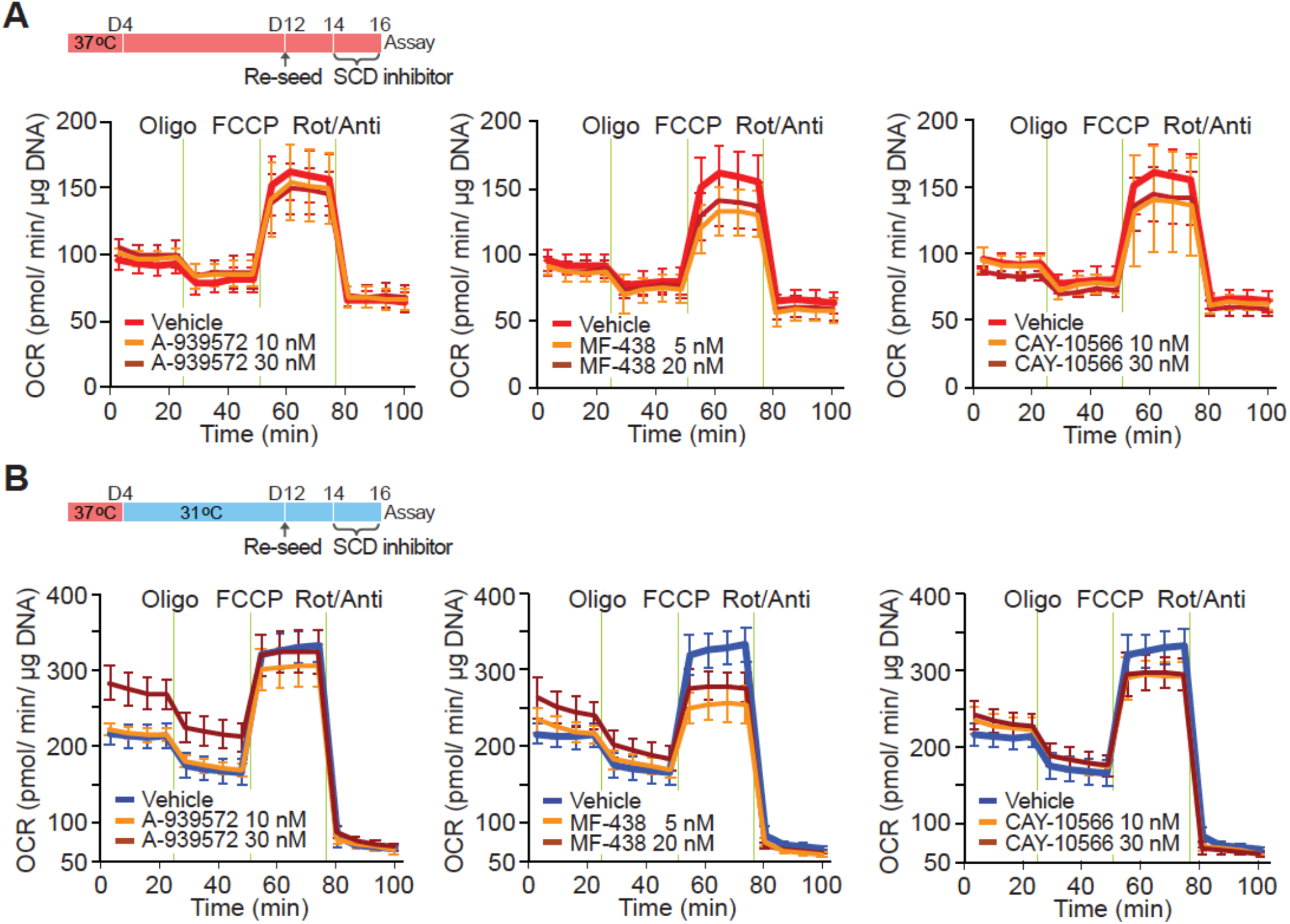
Differentiated adipocytes cultured at either 37°C or 31°C for 8 days were trypsinized, then collected floating adipocytes following centrifugation. Adipocytes were seeded on Cell-Tak coated Seahorse XF96 Cell Culture Microplate, then cultured overnight upside down in order to attach to the plate at either 37°C or 31°C. Adipocytes were cultured with SCD inhibitors: A-939572, MF-438, or CAY10566 at the indicated temperature for two days before the assay. (**A**) No effect of SCD inhibitors on SRC of adipocytes cultured at 37°C (*n* = 8-16 per group). (**B**) Increased SRC of adipocytes adapted to 31°C was inhibited with SCD inhibitors; A-939572, MF-438, or CAY10566 (*n* = 8-16 per group).

